# Proteomics analysis reveals novel phosphorylated residues and associated proteins of the polyomavirus DNA replication initiation complex

**DOI:** 10.1101/2024.02.08.579500

**Authors:** Rama Dey-Rao, Shichen Shen, Jun Qu, Thomas Melendy

**Affiliations:** Department of Microbiology & Immunology, Jacobs School of Medicine & Biomedical Sciences, University at Buffalo, Buffalo, NY, United States of America; Department of Pharmaceutical Sciences, University at Buffalo, State University of New York at Buffalo, Buffalo, NY, United States of America

**Keywords:** DNA replication, SV40, LT, Polprim, RPA, Phosphorylation, Proteomics, Bioinformatics

## Abstract

Polyomavirus (**PyV**) Large T-antigen (**LT**) is the major viral regulatory protein that targets numerous cellular factors/pathways: tumor suppressors, cell cycle regulators, transcription and chromatin regulators, as well as other factors for viral replication. LT directly recruits the cellular replication factors involved in LT’s recognition of the viral origin, origin unwinding, and primer synthesis which is carried out by mutual interactions between LT, DNA polymerase alpha-primase (**Polprim**), and single strand (ss) DNA binding replication protein A (**RPA**). The activities as well as interactions of these three with each other as well as other factors, are known to be modulated by post-translational modifications (PTMs); however, modern high-sensitivity proteomic analyses of the PTMs as well as proteins associated with the three have been lacking. Elution from immunoprecipitation (IP) of the three factors were subjected to high-resolution liquid chromatography tandem mass spectrometry (LC-MS/MS). We identified 479 novel phosphorylated amino acid residues (PAARs) on the three factors: 82 PAARs on SV40 LT, 305 on the Polprim heterotetrametric complex and 92 on the RPA heterotrimeric complex. LC-MS/MS analysis also identified proteins that co-immunoprecipitated (coIP-ed) with the three factors that were not previously reported: 374 with LT, 453 with Polprim and 183 with RPA. We used a bioinformatic-based approach to analyze the proteomics data and demonstrate a highly significant “enrichment” of transcription-related process associated uniquely with LT, consistent with its role as a transcriptional regulator, as opposed to Polprim and RPA associated proteins which showed no such enrichment. The most significant cell cycle related network was regulated by ETS proto-oncogene 1 (ETS1), indicating its involvement in regulatory control of DNA replication, repair, and metabolism. The interaction between LT and ETS1 is validated and shown to be independent of nucleic acids. One of the novel phosphorylated aa residues detected on LT from this study, has been demonstrated by us to affect DNA replication activities of SV40 Large T-antigen. Our data provide substantial additional novel information on PAARs, and proteins associated with PyV LT, and the cellular Polprim-, RPA- complexes which will benefit research in DNA replication, transformation, transcription, and other viral and host cellular processes.

## 1. INTRODUCTION

The polyomavirus (PyV) uses a major viral regulatory protein, large T-antigen (**LT)** that promotes viral proliferation (1). LT is also involved in regulation of timing of the infection cycle, repressing transcription of viral early genes, stimulating expression of viral capsid proteins, and altering transcription of many cellular genes thus preparing the host cells for substantial replication of the infecting virus (2-4). It does that by binding to crucial host cell cycle regulators and tumor suppressors (5, 6) as well as activates the transcription of viral late mRNA (7, 8) and hinders histone deacetylation, leading to an overall activation of transcription (9). LT is the only viral protein required for PyV genome replication, being the origin-binding protein and helicase, as well as recruiting the cellular factors required for replication of the viral DNA. All these interactions make LT an interesting target/tool in the search for compounds with antiviral and/or antiproliferative activities designed for the management of polyomavirus-associated diseases (10, 11).

A schematic depiction of the elongating PyV DNA replication fork (Figure 1) shows LT, which after recognizing the SV40 origin, acts as the hexameric AAA+ DNA helicase and recruits the DNA polymerase alpha-primase (**Polprim**) complex (a heterotetramer of 180, 68, 58 and 48 kDa) and the single-strand DNA binding Replication protein A (**RPA**) complex (a heterotrimer of 70, 32, and 14 kDa), to drive replication of SV40 genomes (2-4, 12-22). All other factors required for initiation, elongation and termination of SV40 DNA replication are provided by the host. It is well known that mutual interactions between LT, Polprim-, and RPA- complexes are critical for PyV DNA replication to be initiated: in origin binding, unwinding the origin of replication, helicase interaction with RPA, loading RPA onto ssDNA, as well as for RNA-DNA primer synthesis (18, 23-33).The clamp loader/sliding clamp (RFC/PCNA) then recruit DNA polymerase delta to carry out processive synthesis of both strands (22, 34, 35). Interactions of LT, Polprim, and RPA are known to be modulated by highly regulated and evolutionarily conserved PTMs that play a pivotal role in DNA replication, DNA repair, paused replication fork stabilization, or other yet unelucidated functions (36-39). Although SV40 is the best understood eukaryotic DNA replication system (40-42), more remains to be understood about the subtleties and regulation of this process, including the inhibition of SV40 DNA replication *in trans* that occurs after treatment of cells with DNA damaging agents (43-47). Elucidating details of PyV DNA replication and its regulation may provide novel avenues to inhibit PyV DNA replication, potentially leading to PyV antivirals for use against progressive multifocal leukoencephalopathy and kidney nephropathy in immunocompromised hosts (10, 11, 48).

**Figure 1.**
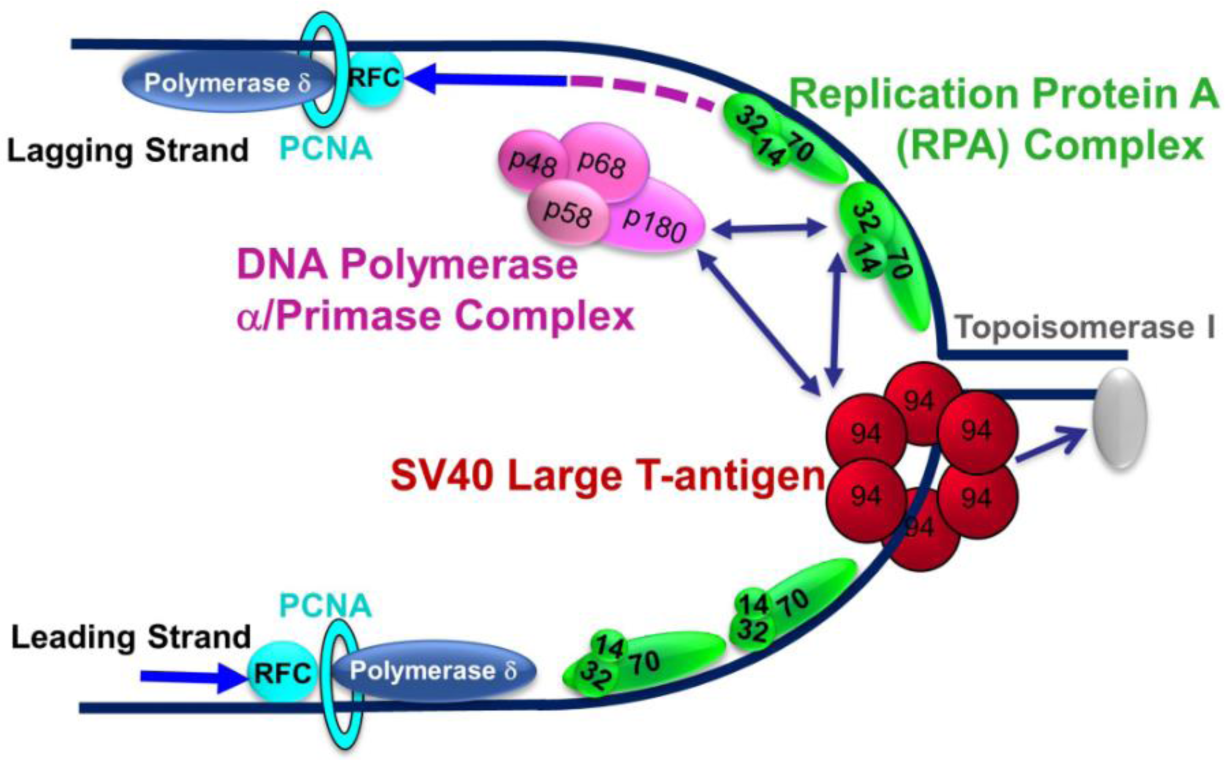
Schematic model of the elongating PyV DNA Replication Fork. Large T-antigen (LT) (monomer ∼90 kDa; red) recognizes the origin of replication and after parking at the origin acts like a helicase to begin unwinding the DNA (dark blue solid line) using ATP hydrolysis for energy. LT recruits the remainder of replication factors required for initiation and synthesis to drive replication of SV40 genomes from the host mammalian cell. The helicase along with Polprim (4 subunits, pink) and RPA (3 subunits, green) interact to begin the process of dsDNA unwinding at the fork and synthesis of RNA/DNA primers (dark pink dashed line on the lagging strand), the critical first steps in PyV DNA replication (bright blue arrows showing directionality). DNA Polymerase delta (blue), recruited through RFC/PCNA (teal) then replicates both the leading and lagging strands. DNA Topoisomerase 1 (TOP1, grey) acts ahead of the fork to relieve super helical tension that would hinder the unwinding of the parental DNA strands.

The paucity of highly sensitive proteomic studies on LT, Polprim- and RPA-complexes has led us to apply immunoprecipitation (IP) combined with mass spectrometry (LC-MS/MS) technology (IPMS) (49) to identify phosphorylated amino acid residues (PAARs) and associated proteins for each of them. We identified 602 PAARs for the 8 polypeptides of LT, Polprim and RPA, of which 479 had not previously been identified. Using phosphomimetic mutation of one of the novel phosphorylated aa residues detected on LT in this study, we have demonstrated a dramatic decrease in DNA replication functions of SV40 Large T-antigen both *in vitro* and in cell culture (50).

An *in-silico* bioinformatic-based approach was used to analyze the co-immunoprecipitated (coIP-ed) proteins, which revealed an enriched <u>transcription-related</u> process network within the SV40 LT associated proteins, notably distinguishing it from Polprim and RPA, and consistent with the known role of LT as a major transcriptional regulator. The 711 non-redundant novel protein associations with either of the three complex clusters formed a “DNA replication signature”, that included common molecules, networks and biological processes previously linked to SV40 DNA replication, validating our data. The transcription factor (TF), TP53 was the most overconnected reaction hub in our dataset mirroring its regulatory significance to the replication-related network. Although the TF, ETS proto-oncogene 1 (ETS1) was absent (‘hidden”) from our dataset, it was the most significantly overconnected TF to our dataset, indicating a potential role in the regulation of viral and cellular DNA metabolism. Both TP53 (∼53 kDa) and ETS1 (∼55kDa) were confirmed to be co-IPed with antibodies to Large T-antigen from HEK293T cells and the LT association with ETS1 was demonstrated to be independent of DNA and/or RNA

## 2. MATERIALS AND METHODS

### 2.1 Cell-Culture and Lysate for IP

HEK293T cells were grown at 37°C with 5% CO_2_ in Dulbecco’s modified Eagle’s medium (DMEM) (Gibco) supplemented with 10% fetal bovine serum (Atlanta Biologicals) and 1% penicillin-streptomycin (Gibco). To induce DNA damage, 70-90% confluent cells were treated with 50μM etoposide solution (Calbiochem) in DMSO (ETO) or were left untreated with only DMSO (Control or Cont.) for 2 hours at 37⁰C. Cell lysates were prepared using NP40 lysis buffer (ThermoFisher) containing 50mM Tris-Cl (pH 7.4), 250mM NaCl, 1% NP40, 5mM EDTA with protease and phosphatase inhibitors (ThermoFisher, Sigma), on ice, subjected to centrifugation in a microfuge for 15 min at 4°C at 15.9K rcf. Protein levels were quantitated using BCA assay (ThermoFisher), flash frozen in liquid nitrogen and stored at -80⁰C.

### 2.2 Antibodies

Monoclonal antibodies mAbs101 or 419 to large T-antigen (LT), mAbs 1644 or 1645 (anti-DNA Polymerase alpha-: POLA1), mAb9 (anti-70 kDa RPA subunit, RPA1), mAb 20 (anti-32kDa RPA subunit: RPA2), and anti-p53 were generated in our laboratory and purified on protein A Sepharose (GE). Commercial rabbit anti POLA1 polyclonal antibodies were purchased from Abcam, rabbit anti-Actin (ACTB) antibody from Sigma. We used two anti-ETS1 antibodies for our LT-IP experiments. One was a mouse monoclonal anti-ETS1 (NBP2-2216 from Novus) that was unable to recognize the TF in the input lysate but identified it IP lanes, and a rabbit monoclonal (MA5-32732 from Thermofisher Scientific) that recognized the TF both in input lane and as coIP-ed protein. All primary antibodies were used at 1:1000-5000 dilution for Western blotting (WB). Rabbit polyclonal antibody (anti-pSQ/TQ) was purchased from Cell Signaling Technology: WB 1:500 in 3% BSA in TBST (50mM Tris-HCl, 150mM NaCl, 0.1% Triton-X100).

### 2.3 IP for Western Blot

We followed the manufacturer’s protocol with the following modifications: A 50 µl slurry of Dynabeads Protein G (as supplied by Invitrogen, Carlsbad, CA Cat # 10003D) was used to immune-precipitate proteins from ∼500µl ETO and DMSO treated HEK293T cell extracts (1.3 mg/ml protein concentration) bound to ∼ 1.0 µg LT, Polprim and RPA -specific monoclonal antibodies. The proteins were eluted from the beads using NuPAGE LDS reducing sample buffer (ThermoFisher), heated to 90°C, resolved on 4-12%, or 4-20% Novex Tris-glycine gels (ThermoFisher) for 1.5h at 150 Volts. Gels were transferred to a nitrocellulose membrane (ThermoFisher) using iBlot2 gel transfer using template program P0. Membranes were blocked for 1h at room temperature with 5% non-fat dry milk (NFDM) in Tris-buffered saline with 0.1% Triton-X (TBST), washed 3 x 5min with TBST. Appropriate primary antibodies (either 1hr at RT or O/N at 4°C with shaking) were used to probe the membranes, washed 3 x 5min with TBST (TBS + 0.1% Triton X-100) and placed in HRP-linked anti-rabbit or anti-mouse secondary antibody (Abcam) (diluted to 0.5μg/mL in TBST for 1h at RT). Membranes were washed 3 x 5 min with TBST and developed using chemiluminescent substrate (ThermoFisher SuperSignal West) and imaged using a ChemiDoc (Bio-Rad). Image J (NIH) was used to quantify bands.

To examine the effect of nucleic acids on the co-immunoprecipitation of LT and ETS1, cell lysates from HEK293T cells were treated with Benzonase (800U/ml) in the presence of 1mM MgCl_2_ for 2hours at room temperature while shaking. Protein lysates were checked for nucleic acid degradation on agarose gel stained with EtBr before adding anti-LT antibodies to the lysate for immunoprecipitation with Dynabeads Protein G as described above. Proteins were resolved on 4-12% Novex Tris-glycine gels and WB was performed as above.

### 2.4 IP for Mass Spectrometry

LT, Polprim and RPA -specific mAbs (approximately 100 µg of each ligand) were separately bound to 5mg of M-280 Tosyl activated Dynabeads (Invitrogen # 14203) O/N at 37⁰C on a rotating wheel, washed twice in 1X PBS and prepared according to manufacturer’s protocol. Binding reactions were set up with the equivalent of 1mg beads + ligand in PBS mixed with 1.3 mg of total protein (whole extracts of Cont. and ETO treated cells separately) at 3.0 mg/ml final concentration for 4h at 4⁰C with tumbling. The beads were washed with PBS three times and eluted with 0.2% formic acid (FA) 4X at RT. The elution was flash frozen in liquid N2 and dried in a speed-vac. Portions of immuno-precipitations were boiled in reducing SDS loading buffer at 95⁰C for 5 min, the bound proteins resolved using 4-12% Novex Tris-glycine gels and silver-stained (MS compatible) (Thermo Fisher Scientific). Trypsin digestions of the 0.2% FA elution were processed for mass spectrometric (MS) analysis at the Proteomics and Bioanalysis Core (PBC) Facility, New York State. The IP followed by MS based proteomics analysis (51) is termed as IPMS in the rest of the paper. We compared results from in-gel digestions (not shown) as well as digestions of the solvent-eluted sample and the latter resulted in many more peptide identifications and better signal to noise ratios. Our results in this report reflect the latter analyses.

### 2.5 Mass Spectrometry (MS)

Liquid chromatography-tandem mass spectrometry (LC-MS/MS) was carried out on a trapping nano-flow LC-Orbitrap MS system consisting of a Dionex Ultimate 3000 nano LC system, a Dionex Ultimate 3000 gradient micro-LC system with an WPS-3000 autosampler, and an Orbitrap Fusion Lumos mass spectrometer (ThermoFisher Scientific, San Jose, CA). For preparation of lyophilized LT samples, 50μL 0.5% SDS was added to each sample, and samples were sonicated for 30 sec and vortex for 10 min to reconstitute protein. Protein reduction and alkylation was performed sequentially by addition of 2μL 200μM DTT and 4μL 500μM IAM, each with 45-min incubation in a covered thermomixer under 37°C with constant shaking. Protein was then precipitated by two-step addition of 60 and 300μL chilled acetone, incubated at -20°C for 3hr, and centrifuged for 30 min at 18,000 g under 4°C to pellet precipitated protein. Pelleted protein was gently rinsed with 400μL methanol, decanted, and wetted using 45μL 50mM Tris-FA. A volume of 5μL trypsin dissolved in 50mM Tris-FA (0.25μg/μL) was added to each sample, and trypsinization was performed under 37°C overnight (∼16hr) with constant shaking in a covered thermomixer. Trypsinization was terminated by addition of 0.5μL FA, and derived peptide mixture was centrifuged at 18,000 g under 4°C for 30 min, and supernatant was transferred to vials for analysis.

A single injection of derived peptides was analyzed for each sample. A large-i.d. trapping column (300µm ID × 5 mm) was implemented prior to nano LC column (75-μm ID × 100 cm, packed with 3μm Pepmap C18) separation for high-capacity sample loading, matrix component removal, and selective peptide delivery. Mobile phase A and B were 0.1% FA in 2% acetonitrile and 0.1% FA in 88% acetonitrile. The 180-min LC gradient profile was: 4−13% B for 15 min; 13% to 28% B for 110 min; 28% to 44% B for 5 min; 44% to 60% B for 5 min; 60% to 97% B for 1 min, and isocratic at 97% B for 17 min. MS was operated under data-dependent acquisition (DDA) mode, with a maximal duty cycle time of 3 sec. MS1 spectra were acquired in the m/z range 400∼1,500 under 120k resolution with dynamic exclusion settings (60 sec ± 10 ppm). Precursor ions were filtered by quadrupole using a 1-Th wide window and fragmented by high energy C-trap dissociation (HCD) at a normalized collision energy of 35%. MS2 spectra were acquired under 15k resolution in either Orbitrap (OT) or Ion Trap (IT).

### 2.6 Data Analysis

LC-MS raw files were searched by Sequest HT (embedded in Proteome Discoverer v1.4.1.14, ThermoFisher Scientific) against SV40 LT sequence. The search parameters include: 1) Precursor ion mass tolerance: 20 ppm; 2) Fragment ion mass tolerance: 0.02 Da (OT)/0.8 Da (IT); 3) Maximal missed cleavages: 2; 4) Fixed modification: cysteine carbamidomethylation; 5) Dynamic modification: methionine oxidation, peptide N-terminal acetylation (spontaneously occurring protein modifications, labeled as O and A in the protein PTM map), serine/threonine/tyrosine (S/T/Y) phosphorylation (labeled as P in the protein PTM map); 6) Maximal modifications per peptide: 4. The spontaneous O and A modifications of free amine groups are not included in this report. LT, Polprim and RPA sequence coverage and peptide lists were exported from Proteome Discoverer. Results were consistent and stable for tryptic peptides of eluted proteins in solution rather than in-gel digestions. The lists of interacting proteins are referred to by Gene IDs (HGNC approved) encoding them for convenience and downstream bioinformatics-based analyses.

### 2.7 Bioinformatics-based Analyses

Prediction of phosphorylated S, T, Y sites: The Netphos v3.1 server (https://services.healthtech.dtu.dk/service.php?NetPhos-3.1) was used to predict Ser-, Thr-, Tyr- (S, T, Y) phosphorylation sites for the full sequences of SV40 LT, PolPrim- (4 subunits) and RPA- (3 subunits) complex. The complete “native” sequence of each protein/subunit was submitted in FASTA format for best prediction, and all three residues (S, T, Y) were selected for the calculation of phosphorylation potential. The default score of ≥0.5 was used as the cutoff for a positive prediction. The server predicts S, T, or Y phosphorylation sites on eukaryotic proteins using ensembles of neural networks. Both generic and kinase specific predictions are performed. The kinase specific predictions are based on consensus cleavage site for 17 kinases (ATM, CKI, CKII, CaM-II, DNAPK, EGFR, GSK3, INSR, PKA, PKB, PKC, PKG, RSK, SRC, cdc2, cdk5 and p38MAPK) that include two of the three PIKK family: Ataxia telangiectasia mutated serine /threonine kinase (ATM) and DNA-dependent protein kinase catalytic subunit or protein kinase, DNA activated, catalytic subunit (DNAPK or PRKDC) involved in DNA damage response (DDR) kinase phosphorylation of several proteins (52). To search for previous literature for binding proteins to SV40 LT, Polprim- and RPA- complexes we used the MetaCore database (https://portal.genego.com), PhosphoSitePlus (https://www.phosphosite.org/homeAction) as well as PUBMED (https://pubmed.ncbi.nlm.nih.gov/).

### 2.8 Ontology Enrichment Analysis

Our IPMS experiments led to identifying coIP-ed proteins (listed by Gene IDs approved by HUGO Gene Nomenclature Committee (HGNC) with the three major proteins (SV40 LT and Polprim- and RPA- complex at the replication fork. Each protein list (Cont. and ETO) (Gene IDs) was uploaded into the web-based integrative software suite from GeneGO: MetaCore+MetaDrug® version 22.2 build 70900 database(Clarivate) (https://portal.genego.com) and mapped to network objects for Gene Ontology (GO) enrichment analyses (53, 54) with the whole human proteome in the background. Hypergeometric tests (55, 56) were used to calculate statistical significance (as p-values and z-scores); in which the null hypothesis is that no difference exists between the number of genes falling into a given category in the target experimental gene list and the genome. Enrichment analysis consists of matching gene IDs of possible targets for the "common", "similar” and "unique" sets with gene IDs in functional ontologies in MetaCore.

To evaluate relevance in biological processes, each pair (Cont. and ETO) of LT, Polprim, RPA-complex associated proteins, was used in a function-based “comparative enrichment analysis” (enrichment by protein function, network building algorithms). The fundamental assumption is that <u>relative connectivity of a gene mirrors its functional significance</u> for the enriched biological processes. Relative connectivity is calculated as the number of interactions between the experimental genes with the genes on the experimental list normalized to the number of interactions it has with all genes in the human proteome in the MetaCore database (Metabase). We focused on the top 10 over-represented process networks (with p-value ≤ 0.01 and false discovery rate ≤ 0.01), that included at least 4 genes. We then made a closer examination of the enriched transcription-related process network that was clearly able to distinguish SV40 LT associated proteins from Polprim and RPA.

### 2.9 Global Analyses

To get a global understanding of the biological context within the vast amount of proteomic data generated, we removed all redundancies in the paired lists of proteins (Gene IDs) (Cont. and ETO) and made a list of proteins that were associated with any or all the three factors (SV40 LT, Polprim- and RPA- complexes). We label this consolidated non-redundant list of seven hundred and eleven (711) proteins (Gene IDs) that coIP-ed with the three factors as the “DNA Replication Signature.” Each entry was associated with the highest spectral counts (SC) detected among all three coIP-ed lysates, which is generally accepted as a semi-quantitative measure for protein abundance in the IP eluate (cell lysates). HGNC approved gene IDs of these 711 proteins were then imported into MetaCore and mapped to ∼800 network objects (due to inclusion of complexes along with individual entries). This is referred to as the “experimental dataset” and is activated for a bioinformatic-based network analyses with the whole human proteome as the background. Unless specified, we refer to proteins by their HGNC approved gene IDs in this report.

### 2.10 Network analyses

The network-based analyses leverage the manually curated knowledge within the MetaCore database (56). We used different algorithms, to drill down into our experimental dataset of seven hundred and eleven gene IDs (representing coIP-ed proteins with LT, Polprim- and RPA-complex)

#### 2.10.1 DNA Replication

To study the networks in the database that are most related to our experimental dataset, we initially used the keyword “DNA Replication” to execute a search of the knowledge base in MetaCore. The most significant of the nine networks associated with the term “DNA replication” was “Cell Cycle-S Phase”, which we evaluated in detail. We externally added SV40 Large T-antigen (not in the human proteome) and allowed the *single interaction algorithm* (*up- and down-stream*) to retrace the network integrating the viral protein into the prebuilt mammalian Cell Cycle-S Phase network from the database that incorporated our experimental dataset as well.

#### 2.10.2 Transcription Factors

To explore significant transcription factors (TFs) that are overconnected to proteins in our dataset associated with biological processes such as DNA replication and cell cycle, we used the *transcription regulation network algorithm*. Networks were built with a variant of shortest paths algorithm with relative enrichment of our activated experimental dataset combined with a relative saturation of networks related to GO processes such as cell cycle, DNA replication and cell death. The networks were prioritized by number of nodes from input list among all nodes in network, p-value and z-score (56). The algorithm output was a list of 30 networks, named by the linked TFs. We examined the top 5 significant networks, (with the largest number of interconnections within the dataset but not necessarily included in it).

#### 2.10.3 Interactome analysis

To investigate connections with the three prioritized replication fork factors (SV40 LT, Polprim (4 subunits), and RPA (3 subunits)) that were used for the IPMS experiments, an interactome analysis was conducted. Using the three as seed nodes we activated the *“build networks*” algorithm “*expand by one interaction*” (*up- and down-stream*) to create merged biological networks. These networks are built on the fly and unique for our activated experimental dataset (gene content as well as spectral counts) with the human proteome in the background. No low trust interactions were included and while retaining the mi-RNA connections we chose not to show names of interacting compounds and drugs (outside the scope of this report).

## 3 RESULTS AND DISCUSSIONS

### 3.1 PAARs on SV40 LT

To initiate viral DNA replication, LT requires specific contacts with the host Polprim and RPA complexes to carry out primer synthesis at the viral replication fork (24, 25, 57-59) that is regulated by PTMs. As a measure of phosphorylation both in control (untreated/Cont.) or etoposide (ETO) treated cells by the DDR pathways, we used the overall SQ/TQ phosphorylation (a preferred ATM/ATR/DNA-PK target sequence) as a proxy. Immunoblotting for phospho-SQ/TQ of HEK293T cell extracts showed a slight increase in overall SQ/TQ phosphorylation in ETO-treated versus Cont in both whole (extr) or LT IP-depleted (depl) extracts of HEK293T cells, as well as in the eluted proteins from IP reactions (Figure 2A: compare lanes 1, 2 and 5 versus 3, 4 and 6). Silver-staining of IP proteins (lanes 7, 8) demonstrated a ∼90 kDa band that coincided with SV40 LT by Western blotting with specific antibodies (lane 11). We compared the LT band in ETO versus Cont IP lanes (Lanes 5, 6) and observed that while the protein amounts remained similar (by silver stain, lanes 7, 8) SQ/TQ LT phosphorylation was∼1.7-fold higher (Image J) in the ETO-treated as compared to Control (compare lane 5 to 6), indicating increased phosphorylation of LT by Phosphatidylinositol 3-kinase-related kinases (PIKKs) due to DNA damage.

**Figure 2.**
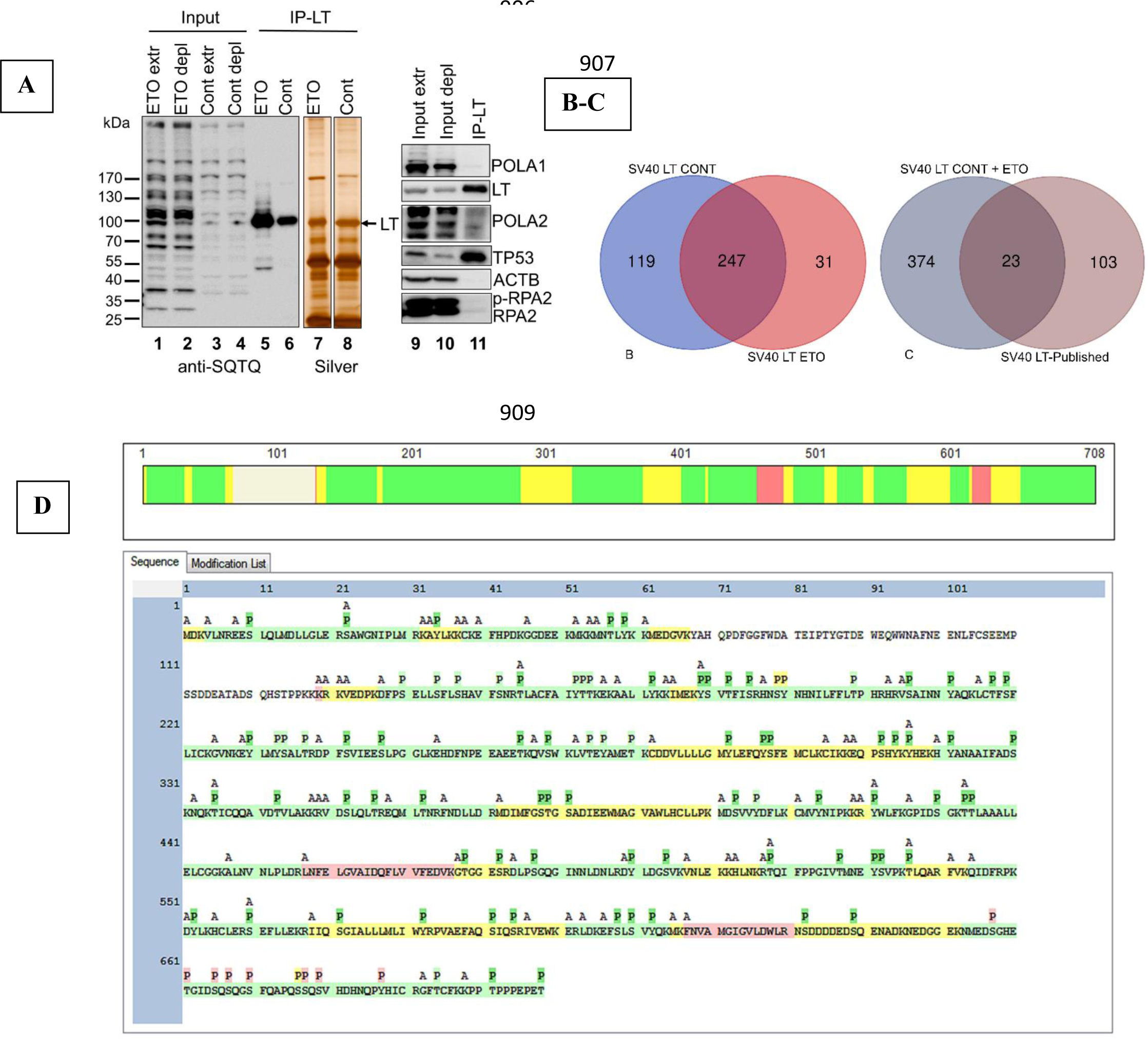
The coIP-ed Proteins and Phospho-map (SV40 LT) IP experiments were performed with SV40 LT-specific monoclonal antibodies from whole cell extracts: untreated (Cont.) and etoposide treated (ETO) HEK293T cells. **(A)** Immunoblotting of whole- (extr) and IP depleted- (depl) cell extracts and IP fractions (lanes 1-6) with anti-SQ/TQ antibodies. The IP eluates were stained with silver (lanes 7, 8). The band at ∼90 kDa coincided with SV40 LT by Western blotting (Lane 11). **(B)** A VENN of the SV40 LT associated proteins for Cont. and ETO shows 247 proteins (gene IDs) in common. For further details of the associated proteins see Table S2. **(C)** We compared the Gene IDs of non-redundant 397 coIP-ed proteins with LT identified by IPMS to the 126 SV40 LT-associated proteins that were previously published to create a VENN diagram of overlap between the two groups. **(D)** Map of phospho-sites on SV40 LT using MS/MS analysis. Only one representative sequence map (with modification) for SV40 LT is shown since there was a large overlap between Cont. and ETO. Sequence coverage of detected peptides (by MS) was 91% for both treatments. Tryptic peptide K67-K129 remained undetected by MS/MS (For further details of SV40 LT PAARs see Table S1). The tryptic peptide detection and S/T/Y phosphor-sites (P) are highlighted with colors by confidence levels (scale bar graphic on top of the full sequence map): green (high) yellow (medium) red (low) and white (undetected by MS/MS). The acetylation of free amine groups is on any aa including Lys (K) and Arg (R) termini and occur spontaneously during experimental steps and may not be biologically meaningful unless validated. To avoid overinterpretation of the data, we do not include acetylated residues in our findings. Abbreviations: P= phosphorylated residues, A= acetylated residues.

Protein phosphorylation is used by cells to transiently change activity of enzymes, interactions, localizations, conformations or even to target them for destruction (38, 60, 61) Here we used high resolution MS2 to catalog the extent of phosphorylation on the three major protein factors in the PyV DNA replication fork. The large number of phosphorylated residues identified in our study is consistent with the abundance of phospho-sites identified on many proteins using advances in mass spectrometric techniques in recent years (38, 62).

A representative sequence map of SV40 LT shows the tryptic peptides (detected by MS) with 82 (Cont) and 85 (ETO) PAARs identified (Figure 2D) out of the total 104 Serine (S), Threonine (T), Tyrosine (Y) that could potentially be phosphorylated. There is a >90% coincidence of PAARs on LT from both treatments, with phosphorylated S378, T379 and S381 found only for ETO but not Cont. A total of 59 S, T and Y residues were predicted to be phosphorylated *in silico* using bioinformatics tools in Netphos v3.1 with the default cutoff score for phosphorylation potential asset at ≥0.5 (Table S1, sheets 1-2). The graphical output is included in Supplementary Figure 1. Our MS2 analysis identified several more phosphorylated S, T, and Ys than the predicted 59 by Netphos; also, there were 8 potential PAARs included on one undetected tryptic peptide in our MS analyses. Although, we found a 1.7-fold higher phosphorylated LT band in ETO compared to Cont by Western blotting, our MS analysis could not quantitate differences between the two treatments. We surmise that since this MS analysis is not quantitative, that even low levels of DDR activation (such as that seen in unperturbed S phase cells) is sufficient to produce detectable phosphorylation on DDR phosphorylation sites in untreated cells. We focused on one of the novel phosphorylated aa residues, T518 detected on LT by this study. Using phosphomimetic mutations on this highly conserved DDR kinase site: we have demonstrated a dramatic decrease in DNA replication activities of SV40 Large T-antigen that uses an alternate method to impede LT helicase activity (50).

The previously reported PAARs on LT were found to cluster into two small regions, one group closer to the amino terminal and the other to the carboxy terminal end of the protein (36, 63-65). It has previously been shown that activation of the LT double hexamer on SV40 origin DNA requires a unique phosphorylation state of LT: modified T124, and unmodified S120 and S123 (64, 66-71). However, these three residues were not identified in this report as they occur in a segment of the protein that was not detected by MS analyses. Of the previously reported 13 phosphorylated aa residues we verified 3 with high confidence (HC) and 4 with low confidence (LC), thus corroborating 54% of the previously published data (70% of those for which we obtained MS data). We add 82 novel PAARs on SV40 LT determined with HC (Table S1, sheets 1-3).

#### 3.1.1 Proteins coIP-ed with SV40 LT

We identified 366 and 278 coIP-ed proteins associated with SV40 LT from Cont and ETO treated extracts respectively (Table S2, sheets 1-3). The Venn diagram (Figure 2B) shows a substantial overlap (247 proteins) common to Cont and ETO, and 119 and 31 unique to each, totaling 397 non-redundant proteins associated with LT. Our work confirmed 23 of the 126 previously reported LT-associated proteins, while discovering 374 additional proteins (Figure 2C). We have thus added a substantial number of novel LT-associated proteins compared to the previous state of knowledge (Table S2, sheets 1-5). Further characterization of the relevance, functional/regulatory significance of the LT-associated proteomics data was accomplished following a bioinformatics-based approach and discussed below.

### 3.2 PAARs on Polprim-Complex

Cell extracts were prepared, subjected to IP for the large catalytic subunit of Polprim, the IP-fractions resolved using SDS-PAGE and stained with silver or immunoblotted for phospho-SQ/TQ, as described above for SV40 LT. As before, an overall difference between levels of phosphorylation at DDR SQ/TQ sites was detected in DNA damaged whole cell- or depleted-extracts (ETO versus Cont). Silver-staining of the IP eluted proteins (Figure 3, Lane 7, 8) revealed almost no differences between Cont and ETO of the ∼180 kDa band identified to be the catalytic subunit POLA1 using Western blotting with protein-specific antibody (Lanes 9, 10). The ratio of this phosphorylated (∼180 kDa) band (compare Lane 6 to 5) was 1.1 (Image J), suggesting that this catalytic subunit of DNA polymerase was not selectively phosphorylated upon DNA damage. IPMS led to identifying all four Polprim subunits in the IP-fractions and like the SV40 LT sequence map described above, only one representative with modifications is provided for each: POLA1(p180, catalytic subunit) and POLA2 (p68, polymerase auxiliary subunit) (Supplementary Figure 2A-B), PRIM1 (p48 primase subunit) and PRIM2 (p58 primase subunit) (Supplementary Figure 2C-D). Results showed a very high coincidence of PAARs between Cont and ETO for each subunit, with a few exceptions (details in Table S3). We detected a total of 184, 79, 62 and 75 phospho- S, T and Ys on POLA1, POLA2, PRIM1 and PRIM2 subunits respectively (Table S3, sheets 1-4) and predicted 124, 81, 35 and 48 phospho-S, T, and Ys *in silico* by Netphos v 3.1 (Table S3, sheet 5) with graphical output (Supplementary Figure 3A-D). For most subunits of the Polprim-complex we detected more PAARs than those predicted *in silico;* however, ∼10-15% of those predicted were not identified by MS, primarily due to the residues being on un- or poorly detected tryptic peptides.

**Figure 3.**
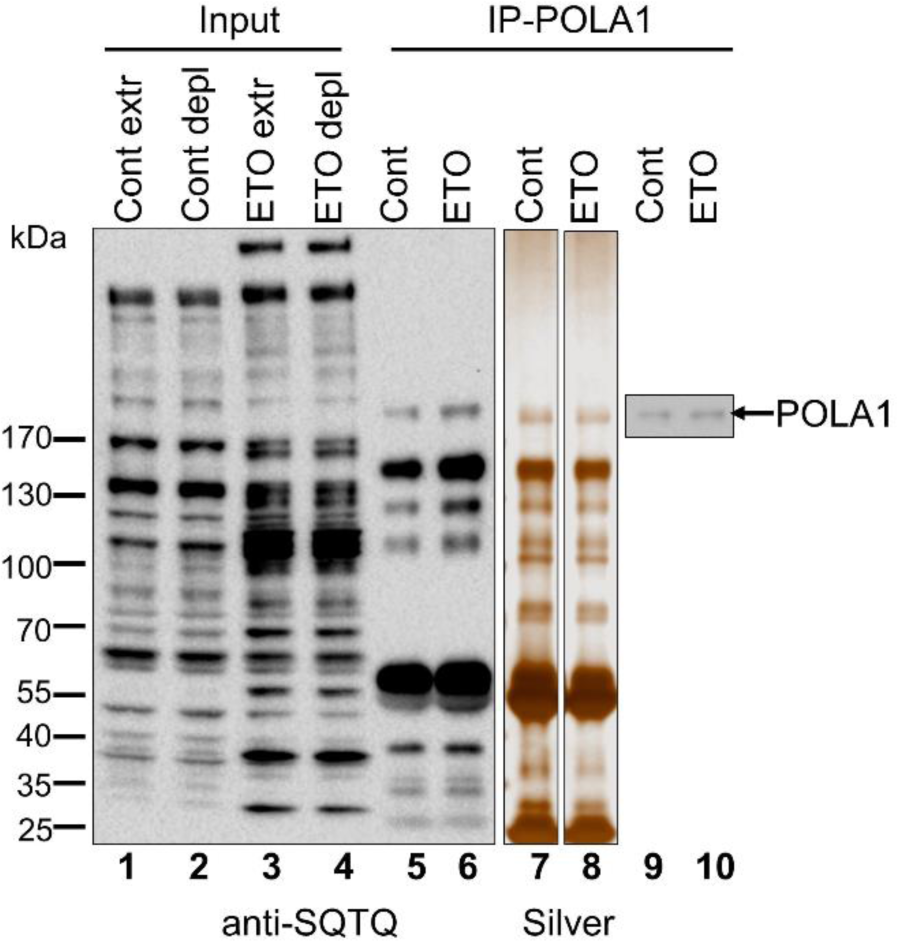
The coIP-ed Proteins with Polprim-complex. The proteins from IP experiments with POLA1-specific monoclonal antibodies were resolved by gel electrophoresis as described. **(A)** Immunoblotting with anti phospho-SQ/TQ antibodies of whole- (extr) and IP depleted- (depl) whole cell lysate and IP fractions from Cont. and ETO treatments (lanes 1-6). IP eluates from Cont. and ETO were stained with silver (lanes 7-8) as well as immunoblotted lane 9-10) and the ∼180 kDa band was identified using rabbit anti-POLA1 antibodies.

Comparing previously published data gleaned from PhosphoSitePlus (72), our analysis was able to confirm all (100%) previously reported phosphorylated residues for POLA1 with HC, 22 out of 24 with HC and two (Y526, Y528) with LC on POLA2 (100%), 13 out of the 15 (87%) with HC and two (S123, S124) with LC on PRIM1 (100%) and finally, 19 out of 20 with HC and one (S492) with LC for PRIM2 (100%), thus corroborating all previously published phospho-data for all subunits of the Polprim-complex. The high level of coincidence and overlap of our experimental results with previously published reports validates our data (Table S3, sheets 6-9). We were able to add 155, 48, 46 and 56 novel phosphorylated S, T, Ys with HC on the respective four subunits of Polprim.

It is well known that Polprim plays a critical role in DNA replication with at least three subunits that interact physically with LT (21). Cell cycle dependent phosphorylation of DNA polymerase alpha inhibits DNA synthesis and may affect Polprim’s physical interactions at the replication fork with either, other replicative proteins or with DNA (73), Polpirm is most active when its regulatory subunit (POLA2) is phosphorylated by CyclinA-Cdk2, and the N-terminal end of its catalytic subunit (POLA1) is not (39, 74, 75). Of the known PAARs on DNA polymerase alpha, the site at the N-terminal end of POLA1, T174, the potential Cdk recognition sites on POLA2, S141, S147, S152 and T156, as well as three putative phospho-sites T115, T127, T130, were confirmed by our work. Access to a larger suite of PAARs on the Polprim-complex than was previously known, will be important to further understand the role it plays in the regulation of DNA replication.

#### 3.2.1 Proteins coIP-ed with Polprim complex

The Venn diagram of coIP-ed proteins with Polprim-complex (Supplementary Figure 4A) shows 144 overlapping between Cont- and ETO-treated cells with 308 and 20 unique to each, respectively (Table S4, sheets 1-3). Our study confirmed 19 out 267 previously reported Polprim-associated factors, while identifying 453 additional ones, thus adding a sizeable number of novel Polprim-associated proteins compared to what was previously known (Supplementary Figure 4B and Table S4, sheets 4-5). The relevance of our findings to DNA replication is investigated using a bioinformatic-based approach and is discussed below.

### 3.3 PAARs on RPA-Complex

As shown above for SV40 LT, there is an increase of overall SQTQ phosphorylation on ETO treatment (DNA damaged) over control (Figure 4 Lanes 1-4). The co-precipitated proteins eluted from beads for both treatments were stained with silver (lanes 7, 8) and immunoblotting identified the prominent silver-stained bands as the 70 kDa RPA1 and 32 kDa RPA2. The slower migrating phosphorylated RPA2 (p-RPA2) is seen in ETO treated IP eluate by SQ/TQ antibodies (lane 6), silver stain (lane 8) and RPA-specific antibodies (lane 13), as well as in ETO treated whole/IP-depleted cell extracts (lanes 11-12) but not in Cont IP fraction (lane 5 & 7), or Cont whole/IP depleted cell extracts (lanes 9-10). While RPA1 and the non-phosphorylated RPA2 band show equivalent levels of phosphorylation in Cont and ETO treatments (ratios close to 1.0 by Image J) (compare lanes 5 and 6), there was a 23-fold increase for the phospho-RPA2 (p-RPA2) band quantitated from ETO versus Cont (compare lane 6 to 5) supporting the known role of RPA2 as being a major phospho-target of the ATM/ATR kinases (43, 76). ATR-dependent phosphorylation is required to inhibit cellular DNA replication in response to DNA damage, although the mechanism remains unknown (77).

**Figure 4.**
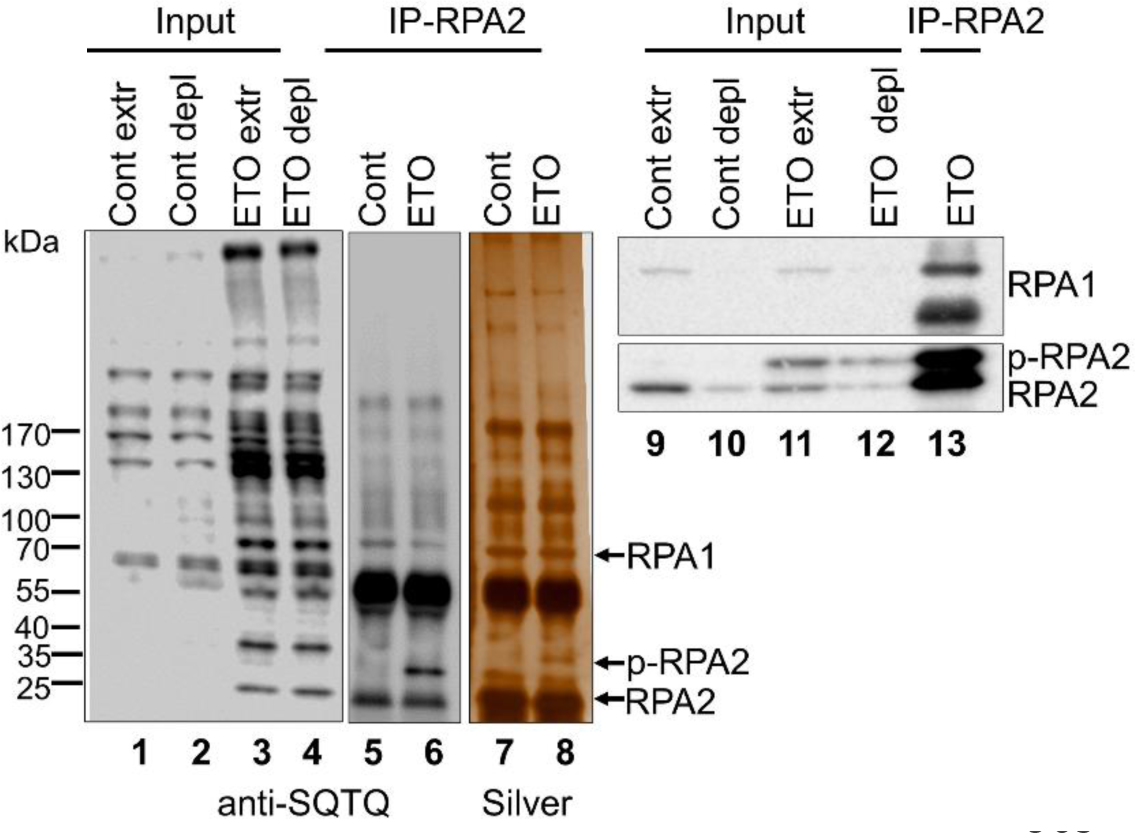
The coIP-ed Proteins with RPA-complex. The proteins from IP experiments with RPA-specific monoclonal antibodies were resolved by gel electrophoresis as before. **(A)** Immunoblot using anti-phospho SQ/TQ antibodies of whole- (extr) and IP depleted- (depl) cell extracts and IP fractions (IP) from Cont. and ETO treatments (Lanes 1-6). The IP elution lanes from Cont. and ETO were stained with silver (Lanes 7, 8). Immunoblot of whole (extr) and depleted (depl) extracts from both Cont. and ETO treatments as well as IP from ETO extract were probed by specific antibodies to POLA1, LT, RPA1, β-Actin (loading control) and RPA2 (Lanes 9-13). The p-RPA2 was clearly identified in the ETO treated samples over control. The three bands (marked with arrows) in Lane 8: 1) RPA1 (70 kDa) 2) unphosphorylated RPA2 (32 kDa) and 3) the slower migrating p-RPA2 were corroborated by Western blotting with specific antibodies (Lane 13).

Our IPMS method led to identifying the three RPA subunits in the IP-eluates using RPA-specific antibodies and a sequence map was generated for each subunit: RPA1 (70 kDa), RPA2 (32kDa) and RPA3 (14kDa) (Supplementary Figure 5A-C) that together bind with high affinity to ssDNA. Sequence coverage for detected tryptic peptides by LC-MS/MS was 100% for all three RPA subunits. We scored the PAARs for each subunit and demonstrated the few differences between Cont and ETO in Table S5, sheets 1-3. It is notable that while we can easily detect the phosphorylated state of the RPA2 (32kDa) band between Control and ETO treated IP by both Western and silver staining (see above) our MS analysis was unable to detect any appreciable differences between them quantitatively. As discussed earlier, this is probably due to the highly sensitive, but non-quantitative LC-MS/MS detection method used. Thus, even infrequent phosphorylation events, such as DDR-mediated phosphorylation events in control cells could be detected. The PAARs identified by IPMS are presented in context of the *in-silico* predictions of 70, 34 and 9 phospho- S, T, and Ys on the 3 subunits respectively via Netphos v3.1 (Table S5, sheet 4) with the graphical output included in Supplementary Figure 6A-C. We discovered a total of 81, 31 and 13 PAARs on RPA1, RPA2 and RPA3 respectively by IPMS. Upon comparing previously published data gleaned from PhosphoSitePlus (72) for the three subunits of the RPA-complex, our analysis was able to confirm 23 out of 24 phosphorylated residues with HC and one (S135) with LC (100%) for RPA1, 6 out of 17 with HC and two (S12 and S33) with LC for RPA2 (47%) and finally 4 out of 4 previously reported phosphorylated amino acid residues for RPA3 (100%). We thus corroborate all previously published phospho-data for both RPA1 and RPA3 and up to 47% for RPA2 (Table S5, sheets 5-7). We were able to corroborate S33 to be phosphorylated on RPA2 albeit with low confidence. S33 phosphorylation was reported previously to be a key event that regulated replication checkpoint brought about by ATR in response to replication stress (78, 79). We add 58, 25 and 9 novel phosphorylated S, T, and Ys with HC on the respective three subunits of the RPA-complex.

The high percentage of previous phospho-data corroborated by this study on all 8 polypeptides of SV40 LT, Polprim and RPA as well as the overall coincidence between our experimental findings and in *silico* predictions, validates our results. The importance and/or relevance of the many phosphorylated residues identified by us on each of the three factors of the replication fork will require future confirmation.

#### 3.3.1 Proteins coIP-ed with RPA Complex

LC-MS/MS analysis of the immunoprecipitated fractions using anti-RPA antibodies identified 225 coIP-ed proteins in both Cont and ETO. The Venn diagram (Supplementary Figure 7A) shows that 43 out of the 45 RPA associated proteins in ETO overlap with Cont There were 180 coIP-ed proteins with Cont and 2 with ETO. (Table S6, sheets 1-3). Our study was able to confirm 42 of the previously reported 673 RPA-associated factors, while identifying 183 novel protein associations (Supplementary Figure 7B and Table S6, sheets 4-5). The low level of similarity of protein associations that we found for all three LT, Polprim and RPA compared to previous literature might be due to the variability in methodologies as well as the transient nature of protein associations/interactions during DNA replication, The large proteomic dataset was further investigated via several bioinformatics-based tools to examine functional relevance and regulatory significance both as paired protein lists from each IP experiment and at a global scale with the trimmed and non-redundant “DNA replication signature”.

### 3.4 Bioinformatics-based Analysis

A function-based comparative analysis of the paired protein lists (Cont and ETO) that coIP with SV40 LT-, Polprim-, RPA- complexes (via MetaCore) revealed the ten most significantly enriched process networks. Among them, “translation elongation termination-” and “cell cycle-” related networks were common to all three comparisons (Figure 5A-C). The SV40-LT associated proteins were uniquely and significantly enriched in a transcription related process network “<u>transcription-mRNA processing</u>” (Figure 5A, lower panel, red box) that included 34 genes common to Cont and ETO,15 unique to Cont and 1 unique to ETO (overall p-value= 2.6E-21, FDR 1.2E-19 included in Table S7, sheet 1; full statistics in Table S7, sheets 1-3). A closer examination of the merged network revealed several TFs (p53/TP53, CDC5L, NRB4/NONO and SFRS10/TRA2B) from our LT-associated dataset and others (PHF5A, TCERG1 and ZNF162) not in our dataset, that connect directly or indirectly to SV40 Large T antigen indicating an involvement of LT with these transcriptional regulators in mRNA processing (Supplementary Figure 8). The evidence is consistent with the unique functionality of the viral protein, LT, as a transcriptional regulator that is capable of recruiting and interacting with the host cellular transcription machinery to regulate several cellular processes including SV40 DNA replication.

**Figure 5.**
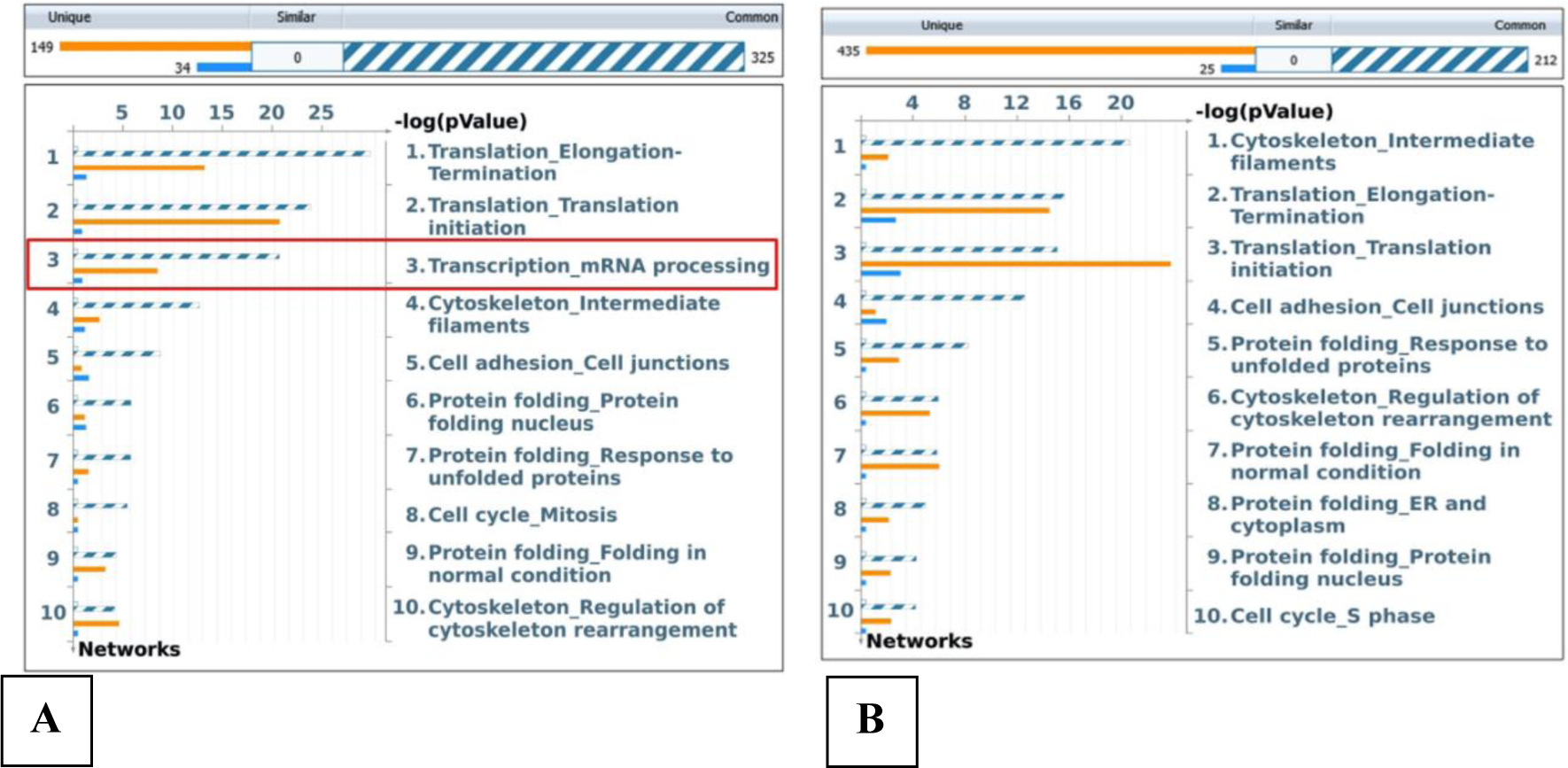
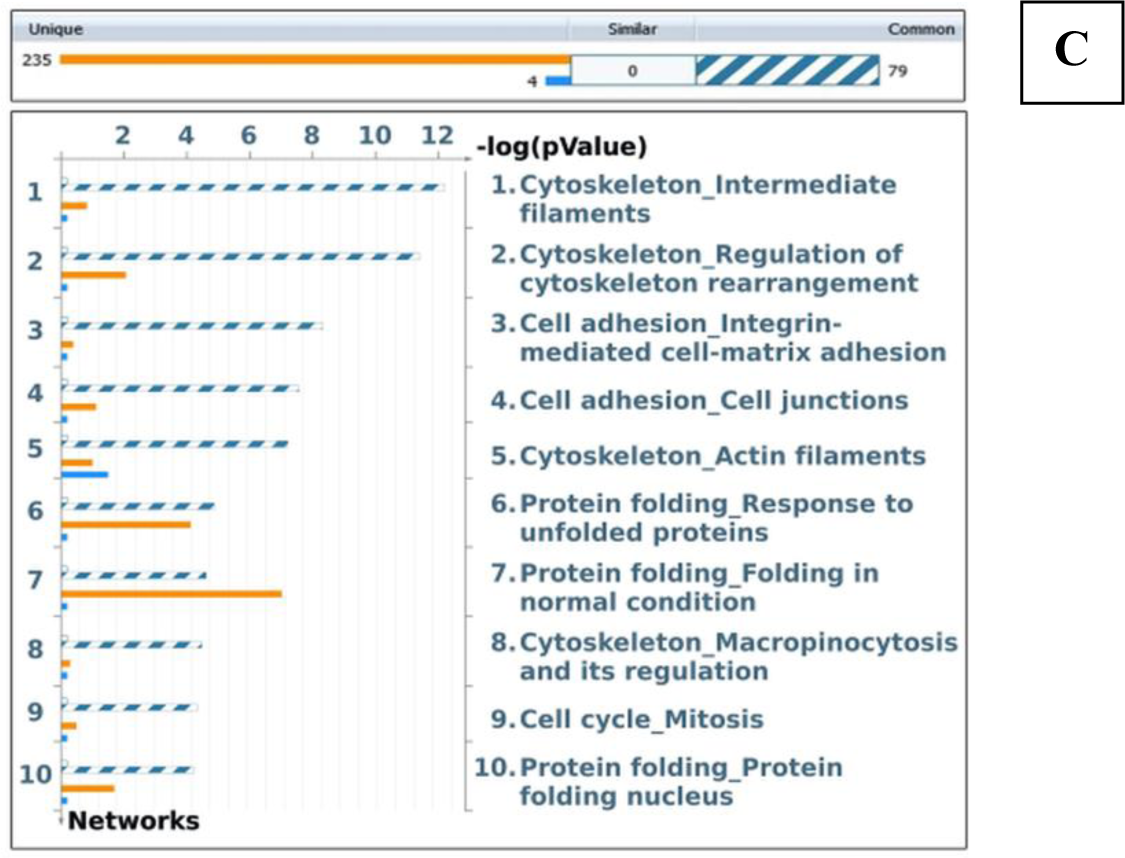
Comparative Enrichment Analysis. The HGNC approved gene IDs (of the coIP-ed proteins with the three factors) were imported into a web-based integrative software suite (MetaCore) and mapped to network objects within the database. The gene content was aligned between the individual uploaded lists (Cont. and ETO) of associated proteins with SV40-LT, Polprim and RPA. The genes sorted into three categories as seen in the histograms: 1) common to both Cont. and ETO (hatched blue and white), 2) unique to Cont. (solid orange) and 3) unique to ETO (solid blue) in the top panels of: **(A)** SV40-LT-associated gene IDs with 325 common, 149 and 34 unique network objects to Control and ETO respectively, **(B)** Polprim-associated gene IDs split into 212 common, 435 and 25 objects unique to Control and ETO respectively and **(C)** coIP-ed proteins (gene IDs) with RPA sorted to 79 common, 235 and 4 unique network objects to Control and ETO respectively. The top 10 enriched process networks are arranged in negative log p-values from most to least relevant in the bottom panels of the three paired analyses (Cont. and ETO) for protein lists from **(A)** SV40 LT, **(B)** Polprim-complex and **(C)** RPA-complex. Sorting is done for p-value of the “common” set. “<u>Transcription-mRNA processing</u>” network was highly enriched and unique in the SV40 LT associated proteins (red box) that distinguished it from those associated with Polprim and RPA-complex. (Statistics: Table S7, sheets 1-3).

### 3.5 Creating a “DNA Replication Signature”

Removing redundancies and merging coIP-ed proteins with LT, Polprim and RPA allowed a global investigation into the relevance of our data to initiation of SV40 DNA replication within the mammalian host cell. A VENN diagram of all coIP-ed proteins in the replication fork revealed the 711 non-redundant proteins/Gene IDs that we termed the “DNA Replication Signature” (Figure 6A-B). Details of the signature (represented by HGNC approved Gene IDs) along with highest spectral counts (a semi-quantitative measure for protein abundance) in each IP reaction are available in Supplementary Table 8, sheets 1-2. The signature showed a considerable overlap (∼42%) with previously reported host protein factors (along with viral LT) known to be associated with the SV40 DNA replication fork (Supplementary Table 8, sheet 3; as well as Figure 1), validating our results to be relevant to “replication fork”, “DNA replication” and “Cell cycle-S phase”. Enrichment analysis by protein function revealed three significant categories (Enzymes, Kinases, and Transcription Factors) with low p-values (6.391E-34, 0.01381 and 0.01014 respectively) and high z-scores (14.65, 2.506 and 2.56 respectively) that are the most overconnected out of several functional categories (Supplementary Figure 9) supported by detailed connectivity statistics (Table S8, sheets 4-5). Several network and interactome analyses were undertaken to reveal functional significance and regulatory relevance of the proteins included within the “DNA Replication Signature”.

**Figure 6.**
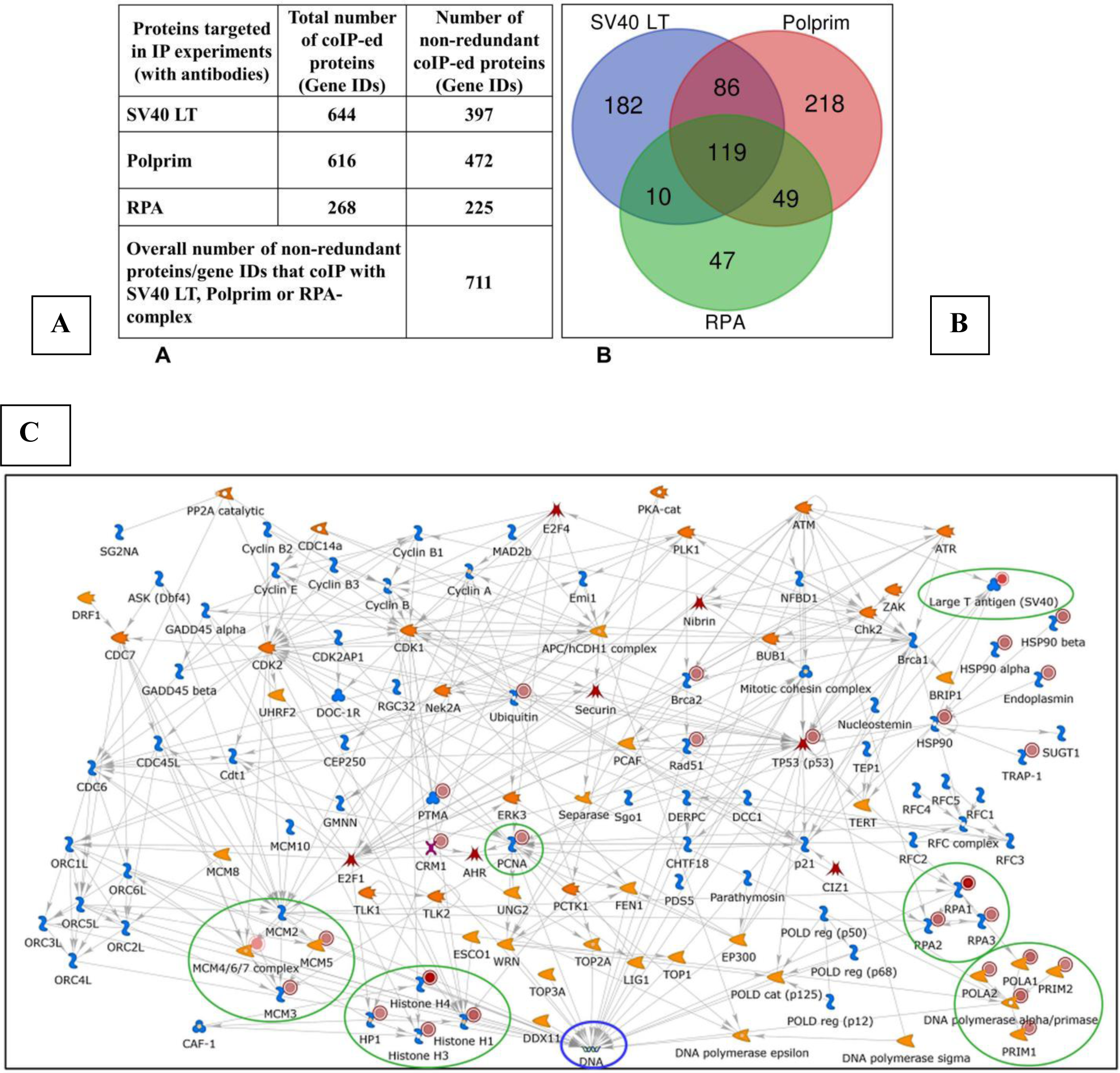
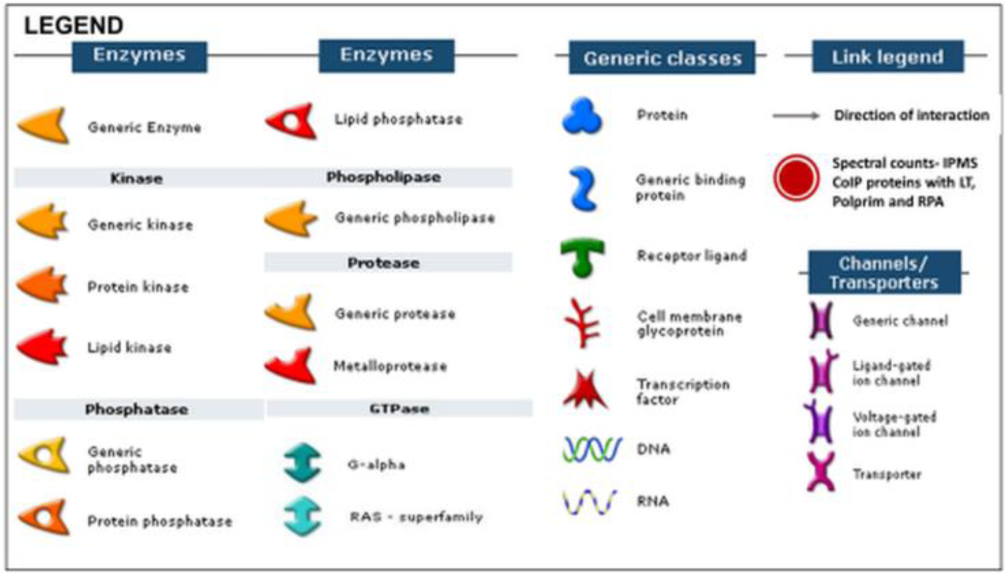
Global analyses. **(A)** Seven hundred and eleven non-redundant proteins (gene IDs) were found to coIP with the three factors needed for initiation of PyV DNA replication: LT, Polprim- and RPA- complex. We call it the “DNA replication signature”. **(B)** A VENN diagram of the 711 gene IDs shows the distribution and overlap between the three factors: 119 of them were shared between all three, with 86, 49 and 10 shared between LT-Polprim, Polprim-RPA and RPA-LT respectively and 182 were unique to LT, 218 to Polprim and 47 to RPA. **(C)** Network Analysis: Cell Cycle_S Phase was the top overconnected network associated with the search term “DNA replication” in the MetaCore database. Many genes from our DNA replication signature were observed within this network (solid red circles) that included a subset known to be associated with the SV40 replication fork (green ovals). Individual proteins or objects are represented as nodes and different shapes of the nodes represent functional classes of proteins and are clarified in the **legend**. (Statistics: Table S9, sheets 1-9). **General information for all networks**: The legend associated with Figure 6C is applicable to all networks. While the enrichments and networks building statistics are calculated according to our uploaded experimental dataset (gene IDs) without spectral counts (SC), the SC are used to set thresholds to allow visualization of the semi-quantitative measure for protein abundance in the IP eluates or cell lysate (solid red circles). The network of interactions (also called edges: grey arrows) between proteins (with direction) were based on the curated knowledgebase within Metacore. The arrowheads indicate the direction of the interactions. Large T-antigen was the only SV40 specific protein added to every network analysis. It mapped to a generic protein in Metacore, to which the red solid circle was added, the color consistent with spectral counts in our MS results.

### 3.6 “DNA Replication” related Network

“Cell cycle_ S phase” was the most significant of the nine networks associated with the term “DNA replication” in the Metabase, that is well documented (74, 80-82). The final merged network (Figure 6C with legend) combined the prebuilt network in the database, with our activated experimental dataset as well as externally added SV40-LT (viral protein that mapped to a generic protein descriptor). Twenty-six gene IDs from our IPMS dataset: LT, PCNA, MCM3, MCM4, MCM5, MCM7, TP53, Rad51, CRM1(XPO1), BRCA2, Ubiquitin, HSP90 family: HSP90 alpha (HSP90AA1), HSP90 beta (HSP90B1), PTMA, TRAP1 and members of the Histone family, subunits of Polprim- and RPA-complex, were included among the Gene IDs that were included in this network. Several among them were also previously reported to be associated with the SV40 DNA replication fork (Table S8 sheet 3). The network has 40 interaction hubs (Gene IDs with 6 to 31 connections) with DNA as the hub where maximum reactions converge (convergence hub). Of the seven TFs included in the network only one, TP53 (with 28 interactions) was from within our dataset, the rest are not (thus considered “hidden”). The significant biological processes enriched within the network were related to DNA metabolic process, DNA replication, cell cycle, and DNA damage and repair, among others and included several genes from our results (Supplementary Figure 10). Several proteins such as MCM3, MCM5 MCM4 MCM7 that are part of the mammalian replication machinery were found to coIP with SV40 LT and Polprim. In summation, this network validates our IPMS protein dataset to be representative of both SV40 and human DNA replication in the cell cycle. The enrichments and network building statistics were calculated according to our experimental dataset (For statistics underlying the “Cell cycle_ S phase” network: Table S9, sheets 1-9).

### 3.7 Transcriptional Regulation Network

The most significantly overconnected network using the “transcriptional regulation network” algorithm was named by the associated TF, ETS1 (p-value =0, z Score = 524.25) (Table S10 Sheet 2). The merged network (with externally added LT) was connected to ∼118 genes from within our dataset. (Figure 7A) and associated with cell cycle and apoptotic process among others (Table S10, sheet 11). We discovered that while ETS1 (an interaction hub with 106 connections) itself was not included within our experimental dataset, out of the thirty-three interaction hubs (each with more than 5 connections) in the network, TFs such as TP53 (with 55 interactions) and TBX2 (with 54 interactions) were from within our DNA replication signature. For details and statistics underlying the network see Table S10, sheets 1-11. The overconnectivity of TP53, and TBX2 from within our dataset and ETS1 hidden from it demonstrates the likely importance of all three transcription regulators in SV40 and/or mammalian DNA replication.

**Figure 7.**
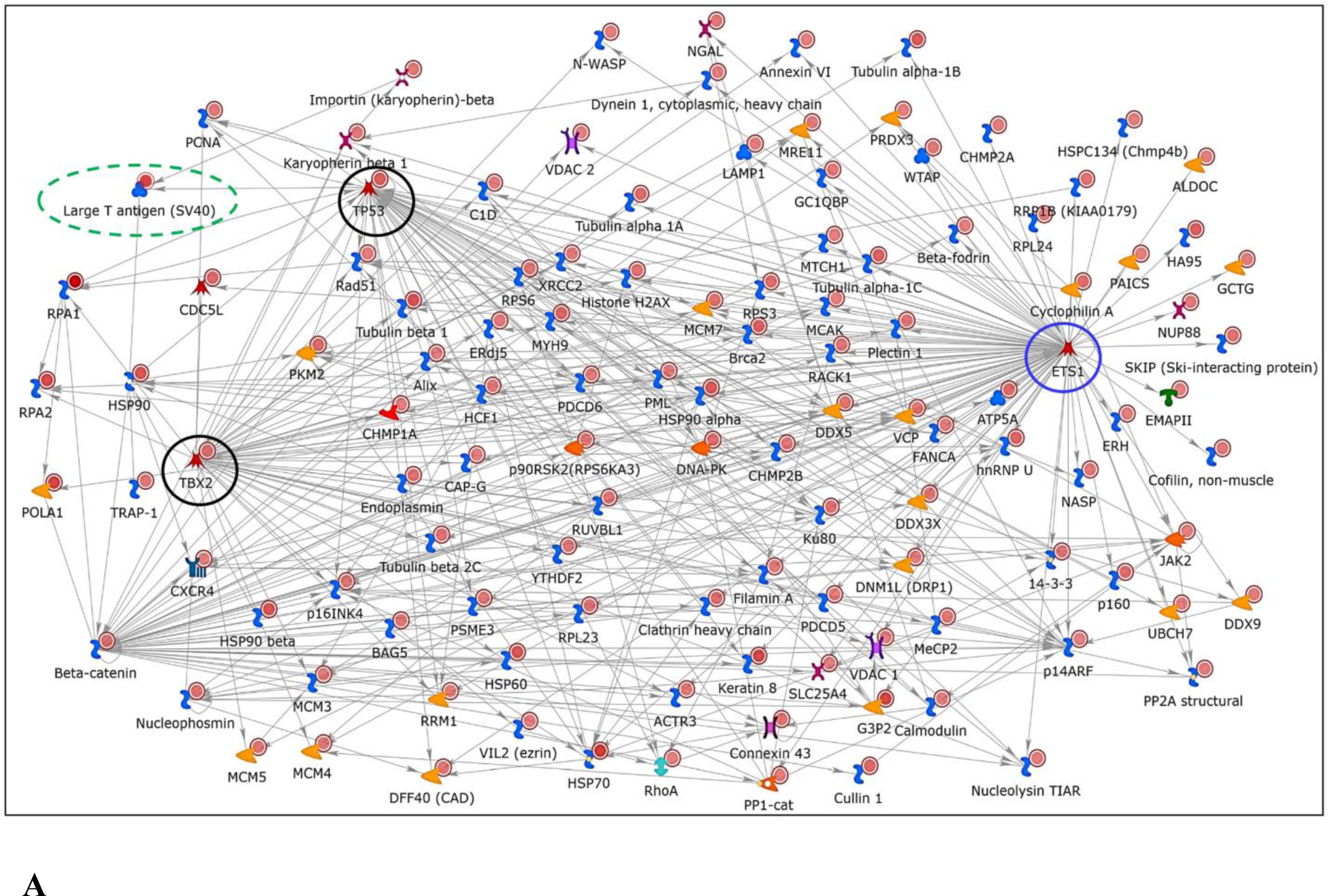
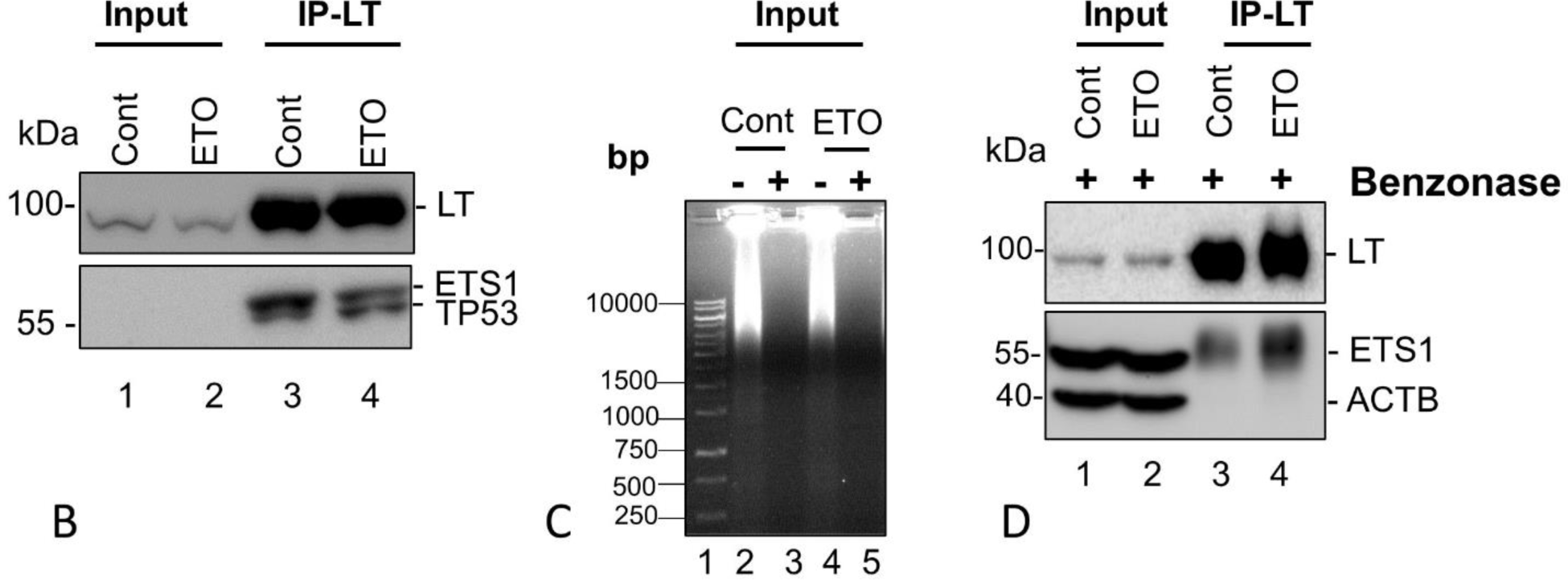
Transcription Regulation Network finds a “hidden” transcriptional factor ETS1. **A)** Using the “transcriptional regulation network” algorithm we generated a list of the top 30 networks named by the most overconnected transcriptional factors (TFs) to our dataset. We selected the top network, named for its TF (ETS1) and observed the biological network enriched with proteins/genes from within our DNA replication signature (solid red circles). The merged network is redrawn after adding Large T-antigen manually (green oval). ETS1 (blue circle) with 106 interactions is predicted to be the most overconnected transcription factor to our dataset although it was unidentified in our dataset (considered to be “hidden”). Our dataset also showed overconnectivity with TFs such as TP53, TBX2 (black circles) with 55 and 54 interactions respectively. (Statistics: Table S10, sheets 1-11). **B)** Immunoprecipitation experiments were performed with HEK293T whole cell lysates (Cont and ETO- Input) using antibodies to Large T-antigen (LT) as described in Methods. The ETS1-encoded protein, Ets1, co-immunoprecipitated with LT along with TP53/p53 (Lanes 3, 4). To examine the effects of nucleic acids on the association of LT with Ets1, DNA /RNA were degraded in the cell lysates by Benzonase nuclease treatment before co-IP experiments were performed. **C)** proteins were run on an agarose gel stained with ethidium bromide to confirm the breakdown of nucleic acids. **D)** Western blots of the input and LT-IP proteins were probed with anti-Ets1, p53 and β-Actin antibodies. ETS1/Ets1 to be co-immunoprecipitated with LT even after Benzonase treatment.

To determine whether the predicted but “hidden” ETS1 associates with LT we performed a co-IP experiment using anti-LT antibodies with both control and ETO treated whole cell lysates from HEK293T cells. A Western blot (Figure 7B) probed with anti-LT, Ets1 and p53 antibodies clearly identifies the ∼55kDa Ets1 protein that co-IPs with LT and runs very close to TP53/p53 at ∼53kDa (Lanes 3, 4). To ascertain that the association of LT and Ets1 is independent of nucleic acids, we removed DNA and RNA from HEK293T cell lysate (Cont and ETO) with Benzonase-nuclease treatment before the IP reaction (Figure 7C). Ets1 and p53 (not shown) co-immunoprecipitated with LT even after Benzonase treatment. A different anti-Ets1 antibody clearly identified the TF in the input lanes as well as a co-IPed protein (Figure 7D). This novel association of Ets1 with LT reported here for the first time is independent of nucleic acids. ETS proto-oncogene 1, transcription factor (Ets1) is a member of the ETS family of evolutionarily conserved sequence-specific DNA-binding transcriptional activators that have been associated in tumor progression (83, 84) and play important roles in diverse cellular processes such as proliferation, differentiation, lymphoid development, motility, invasion, and angiogenesis that are likely to be dependent on specific protein interactions (83-85). Since SV40 LT is a major transformation-inducting protein and is known to target several tumor suppressors such as tumor protein p53 (TP53), RB transcriptional corepressor 1 (RB1) and mitotic checkpoint serine/threonine kinase (BUB1) among others, our findings could fuel further study of SV40 Large T antigen targeting the transcriptional protooncogene, ETS1 to investigate its specific regulatory function in cellular transformation.

The three major replication fork complexes are known to be connected to one another directly for effective replication initiation (18, 23-33) and were found to coIP with each other in our IP experiments. POLA1, POLA2 and RPA2 coIP-ed with LT, the four subunits of Polprim and three subunits of RPA coIP-ed with Polprim and all three RPA subunits with RPA. All three, used as seed nodes were observed to be connected to one another in an *in silico* <u>interactome analysis</u> (Figure 8) via 2 proteins implicated in replication and cell cycle progression: 1) Histone Deacetylase 3 (HDAC3) known to catalyze the deacetylation of lysine residues of the core histones playing an important role in cell cycle progression (85) and 2) Specificity protein1 (SP1), a transcription factor, known to regulate expression of a large number of genes involved in cell growth (86). The only connector protein observed between LT and Polprim in this network is Sirtuin1 (SIRT1) that was shown to bind with HDAC1 and HDAC3 to deacetylate and activate LT (85) acting as a gatekeeper of replication initiation (87). Additionally, LT is observed to be connected to the RPA-complex (directly or indirectly as both inhibitor and activator) via three genes (TP53, EP300 and ATM) that have been implicated in replication and cell cycle progression. Of note, TP53 gene encoding the TF tumor protein p53 (TP53) contains a transcription activation, DNA binding and oligomerization domain and is a known checkpoint protein in cell cycle as well as one of the LT binding proteins (88) and was shown to be enriched at replication forks and nascent chromatin in HEK293T cells probably stabilized due to E1A and LT expression (89). SV40 LT binding to p53 has been long described (90) and acts as a regulator of checkpoint/apoptosis. It acts as a tumor suppressor and regulates cell division by preventing cells from growing uncontrollably (91, 92). When DNA gets damaged by toxic chemicals, radiation, or UV, it can determine whether the DNA will be repaired, or the cell will die (undergo apoptosis) (91, 93). It is also known to respond to several cellular stresses and regulates processes such as cell cycle arrest, apoptosis, senescence, repair of DNA or even changes in metabolism (94-96) and associates with SV40 LT to bind to E1A binding protein p300 (EP300) to direct the acetylation of SV40 LT (97). ATM serine/threonine kinase (ATM) is a member of the PIKK family involved in the phosphorylation of RPA2 as part of the DDR response that is essential for optimal SV40 replication in primate cells (98-100).

**Figure 8.**
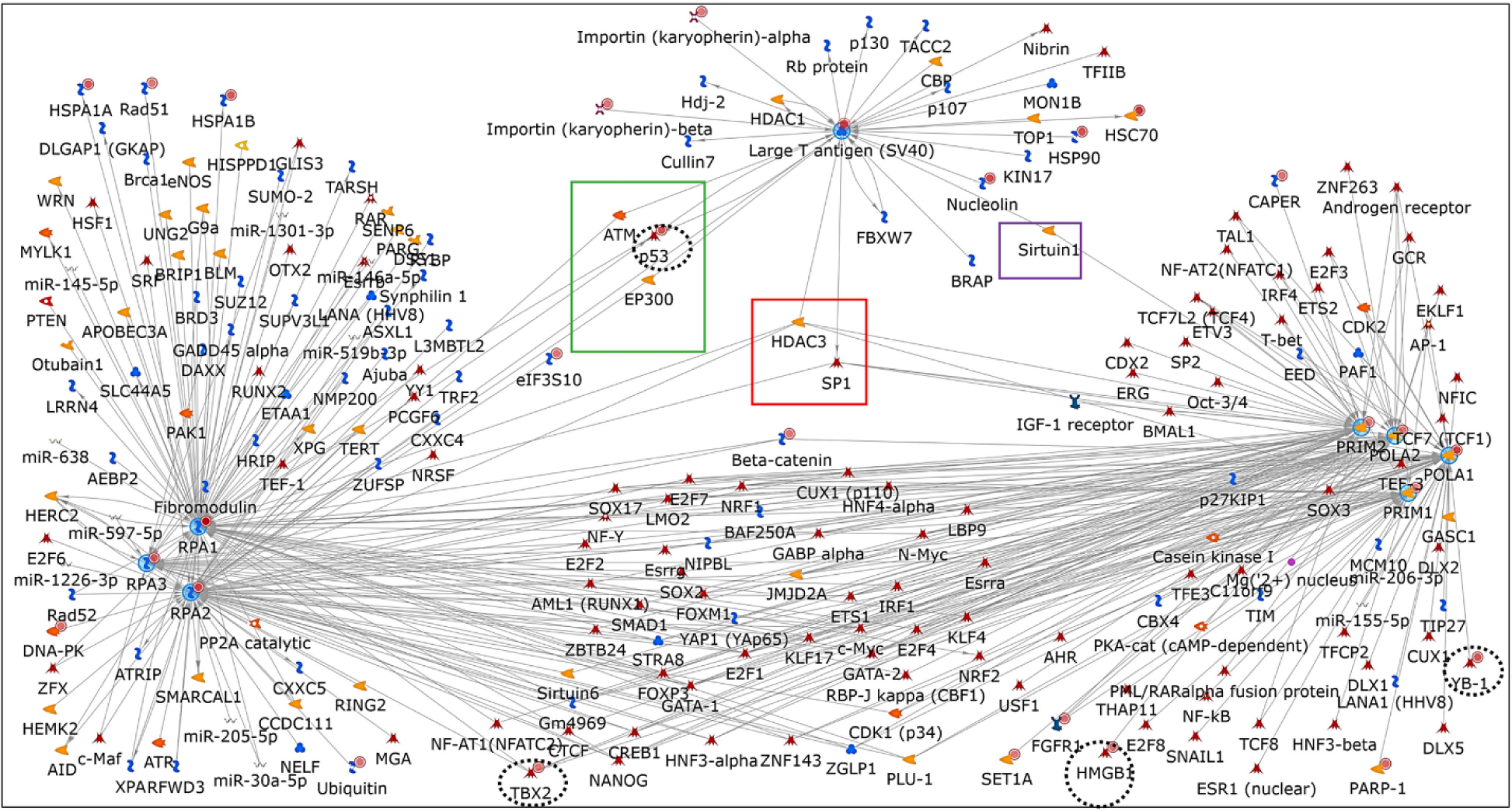
Interactome Analysis. Interaction networks were built around the three protein factors with subunits (seed nodes) (used in our IPMS experiments) acting as hubs (with five or more interactions/edges: grey arrows): SV40 LT, the tetrameric Polprim-complex and the trimeric RPA-complex. LT is linked with Polprim via Sirtuin (purple box). LT is connected to RPA via 3 proteins: ATM, EP300 and TP53 (p53) (green box). All three factors are connected to each other via two proteins: HDAC3 and SP1 (red box). Among the 102 TFs that the three factors interact with, only 4 were from within our dataset (black dashed circle): HMGB1 and YB-1: interact with subunits of the Polprim complex in this network, TBX2 is linked to subunits of RPA- and Polprim- complexes, and TP53 (p53) is the link between RPA and LT. Twenty-seven genes from our activated dataset are included within this network and are associated with solid red circles. Please see “General information for all networks” at the end of Figure 6 description.

Twenty-seven proteins that were included in this interacting network coincide with our experimental dataset. LT was associated with only 30 interactions while RPA-complex (with RPA1: 92, RPA2: 75, RPA3: 29 interactions) and Polprim- complex (with POLA1: 71, POLA2: 54, PRIM1: 35, and PRIM2: 51 interactions) showed much higher interconnectivity to genes (from within or outside our experimental dataset). This is to be expected since LT is a viral protein that can interact with factors within the cell and bring about viral replication but would not be expected to be as interconnected as cellular proteins. While the 3 factors in the replication fork were regulated by ∼102 TFs only 27 were major hubs of interaction. Of them, only TP53 (p53), T-box transcription factor 2 (TBX2), Y-box binding protein 1 (YBX1 or YB-1), and high mobility group box 1 (HMGB1) were from within our data (circled in dashed black). Mapping studies have established that the N-terminal half as well as the ssDNA binding domain of RPA1 are interaction sites for POLA1 and essential for polymerase primer synthesis, while the interaction with LT interfered with the RPA1-POLA1 interaction. The competition between LT and POLA1 for RPA might be playing a crucial role in switching POLA1 function from priming to DNA synthesis (28). Studies such as this help to identify new players in the process and add to the body of knowledge on SV40 DNA replication as an important model for eukaryotic DNA replication.

## 4 SYNOPSIS

Results from this study have provided a plethora of previously unidentified phosphorylated amino acid residues (PAAR) on the key cellular DNA replication/repair/DNA damage response complexes, PolPrim and RPA, as well as the SV40 LT protein, which is involved in these pathways, along with being a major transformation-inducting protein. The novel PAAR sites thus provide new avenues for investigation of regulation of all these important interactions and associations SV40 has with its mammalian host. Drawing on the discoveries from this investigation, we show that phosphomimetic mutation of one of the novel PAAR sites on SV40 LT first identified in this study results in a dramatic decrease of only the DNA replication function of LT, both biochemically (*in vitro*) and in cultured cells (50). In addition, this study has provided hundreds of additional proteins found to be associated with the three factors. Bioinformatics-based analyses of their inter-relatedness are consistent with known functions but have also identified a key interconnected transcription factor, ETS1, that may play a hitherto undetermined more direct role in these processes. We have validated the association between the transcription factors LT and ETS1 and show it to be independent of the presence nucleic acids. Together our results confirm several previously known PAARs, as well as proteins associated with LT, PolPrim-, and RPA-complexes, and have identified numerous more to fuel further investigations.

## Supporting information

PAARs SV40LT

coIP-ed proteins with SV40LT

PAARs Polprim-complex

coIP-ed proteins with Polprim-complex

PAARs RPA-complex

coIP-ed proteins with RPA-complex

Statistics underlying Enrichment Analysis Statistics

DNA Replication Signature

Statistics underlying Cell Cycle S-Phase Network

Statistics underlying ETS1 Transcription Factor Network

Results of Bioinformatic-based Analyses

## AUTHOR CONTRIBUTIONS

Conceptualization and Methodology: RDR and TM; Software and Validation and formal analyses, data curation: RDR and SS; Investigation, Visualization: RDR; Funding Acquisition: TM; Resources: SS and JQ performed mass spectrometric analysis at the Proteomics and Bioanalysis Core (PBC) Facility at University at Buffalo. All bioinformatic-based analyses of Proteomic data using statistical and computational techniques on the MetaCore platform: RDR; Roles/Writing-original draft: RDR; Writing -review and editing: RDR and TM. All authors reviewed and approved the final submitted version of the manuscript.

## DATA AVAILABILITY STATEMENT

The raw data supporting this report is included in the Supplementary Material and all raw data will be made available by the authors without undue reservation.

## SUPPLEMENTARY MATERIAL

The Supplementary material supporting this article can be found online at:

