## Supplementary material for "Proteomics analysis reveals novel phosphorylated residues and associated proteins of the polyomavirus DNA replication initiation complex": Results of Bioinformatic-based Analyses

### Slide 1
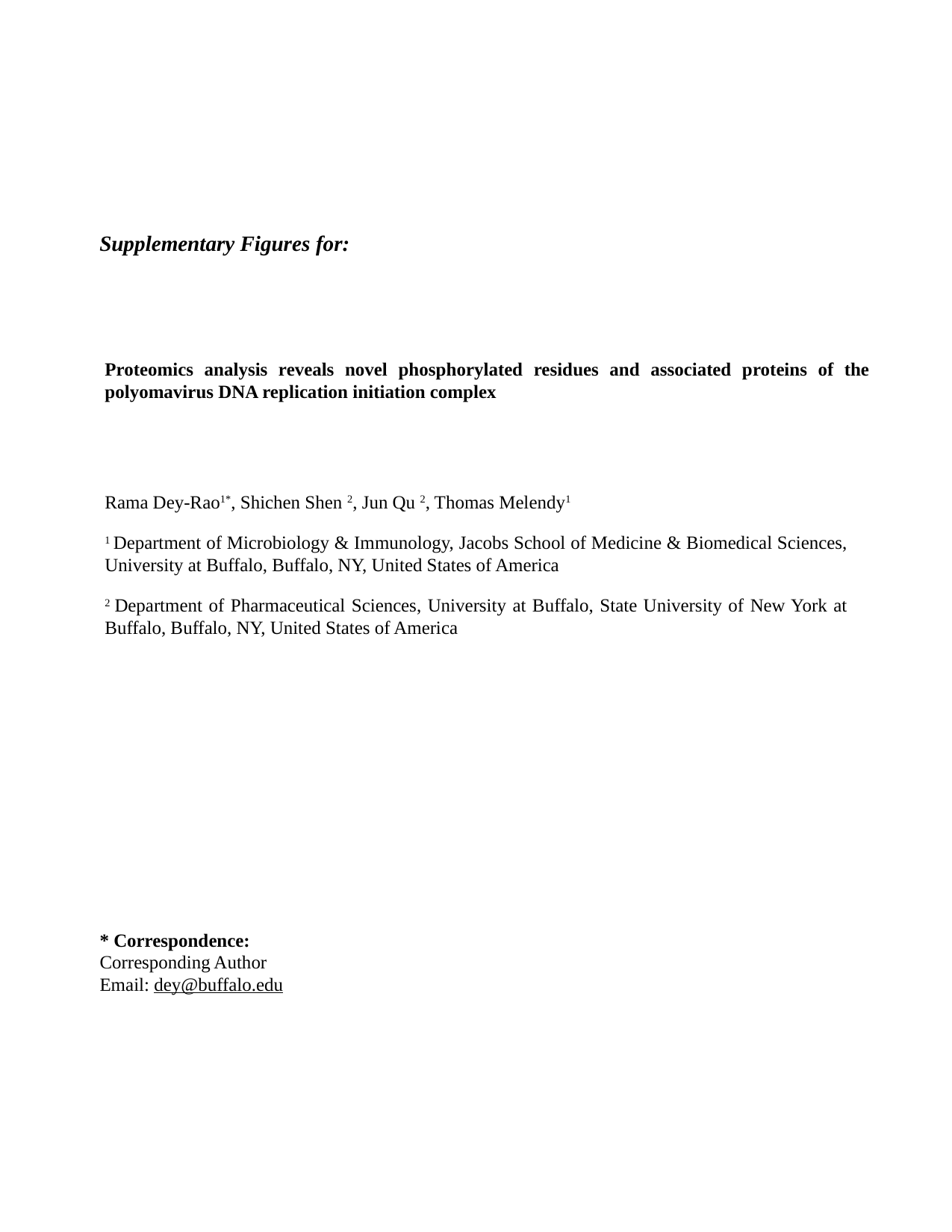

Supplementary Figures for:
Proteomics analysis reveals novel phosphorylated residues and associated proteins of the polyomavirus DNA replication initiation complex
Rama Dey-Rao^1*^, Shichen Shen ^2^, Jun Qu ^2^, Thomas Melendy^1^
^1^ Department of Microbiology & Immunology, Jacobs School of Medicine & Biomedical Sciences, University at Buffalo, Buffalo, NY, United States of America
^2^ Department of Pharmaceutical Sciences, University at Buffalo, State University of New York at Buffalo, Buffalo, NY, United States of America
* Correspondence:
Corresponding Author


### Slide 2
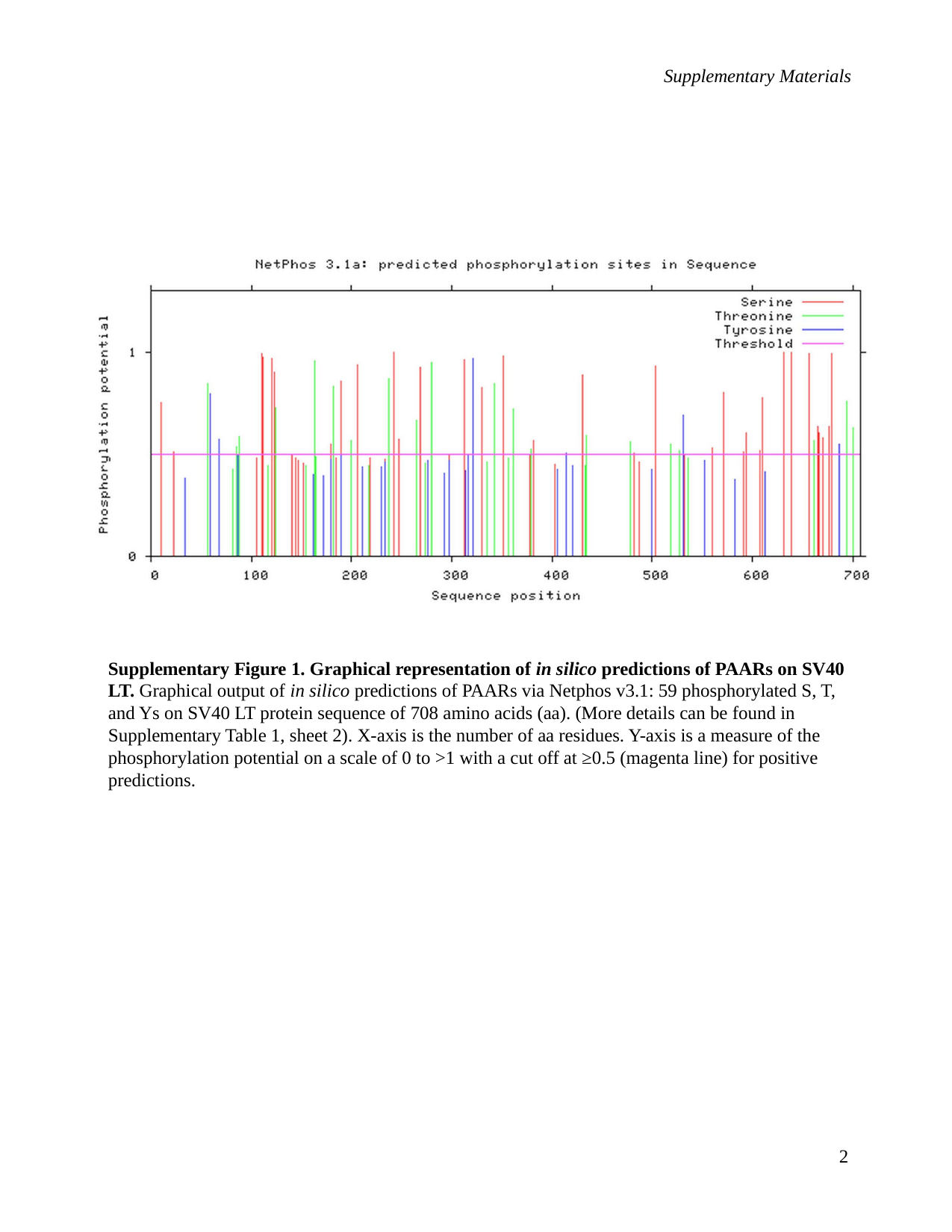

Supplementary Materials
Supplementary Figure 1. Graphical representation of in silico predictions of PAARs on SV40 LT. Graphical output of in silico predictions of PAARs via Netphos v3.1: 59 phosphorylated S, T, and Ys on SV40 LT protein sequence of 708 amino acids (aa). (More details can be found in Supplementary Table 1, sheet 2). X-axis is the number of aa residues. Y-axis is a measure of the phosphorylation potential on a scale of 0 to >1 with a cut off at ≥0.5 (magenta line) for positive predictions.
2

### Slide 3
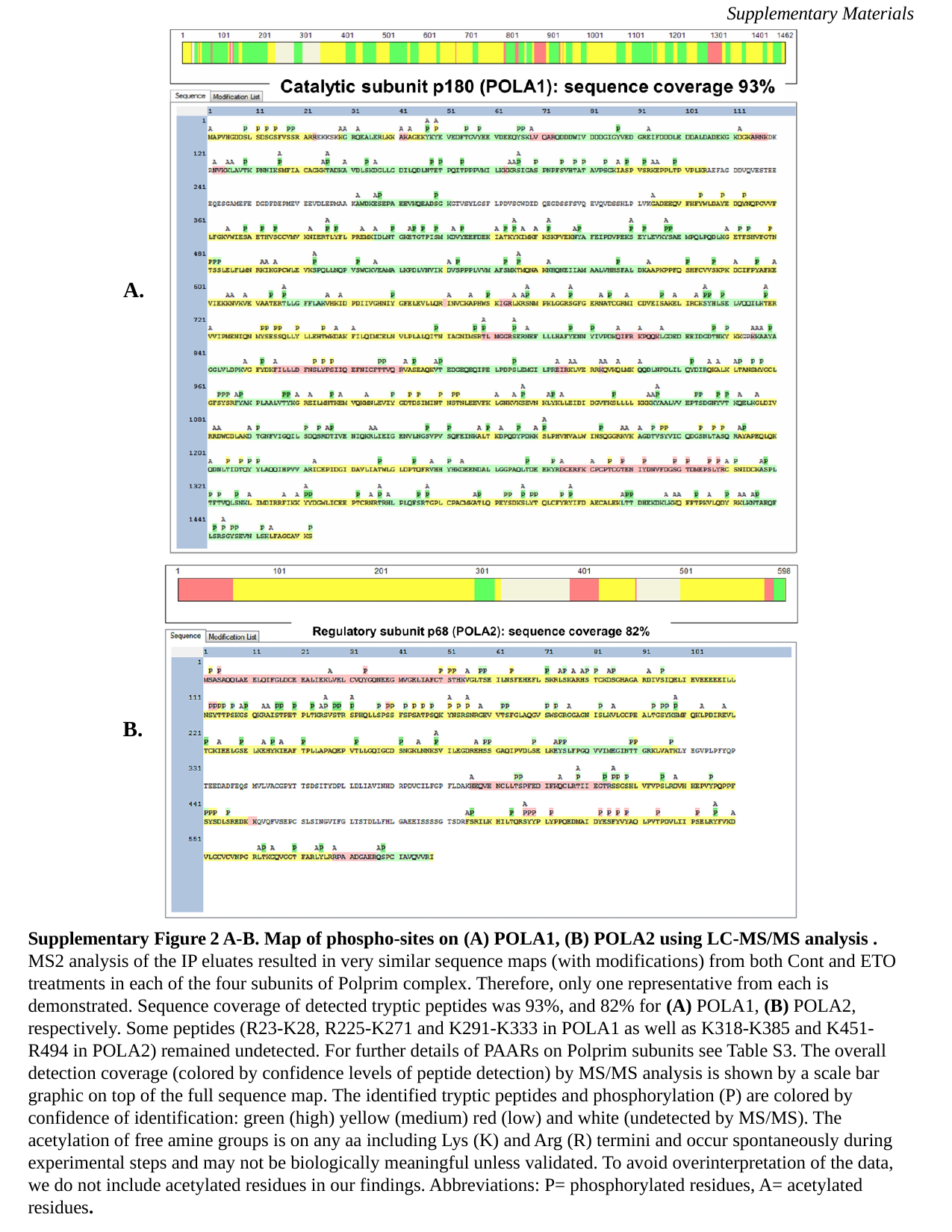

Supplementary Materials
A.
B.
Supplementary Figure 2 A-B. Map of phospho-sites on (A) POLA1, (B) POLA2 using LC-MS/MS analysis . MS2 analysis of the IP eluates resulted in very similar sequence maps (with modifications) from both Cont and ETO treatments in each of the four subunits of Polprim complex. Therefore, only one representative from each is demonstrated. Sequence coverage of detected tryptic peptides was 93%, and 82% for (A) POLA1, (B) POLA2, respectively. Some peptides (R23-K28, R225-K271 and K291-K333 in POLA1 as well as K318-K385 and K451- R494 in POLA2) remained undetected. For further details of PAARs on Polprim subunits see Table S3. The overall detection coverage (colored by confidence levels of peptide detection) by MS/MS analysis is shown by a scale bar graphic on top of the full sequence map. The identified tryptic peptides and phosphorylation (P) are colored by confidence of identification: green (high) yellow (medium) red (low) and white (undetected by MS/MS). The acetylation of free amine groups is on any aa including Lys (K) and Arg (R) termini and occur spontaneously during experimental steps and may not be biologically meaningful unless validated. To avoid overinterpretation of the data, we do not include acetylated residues in our findings. Abbreviations: P= phosphorylated residues, A= acetylated residues.

### Slide 4
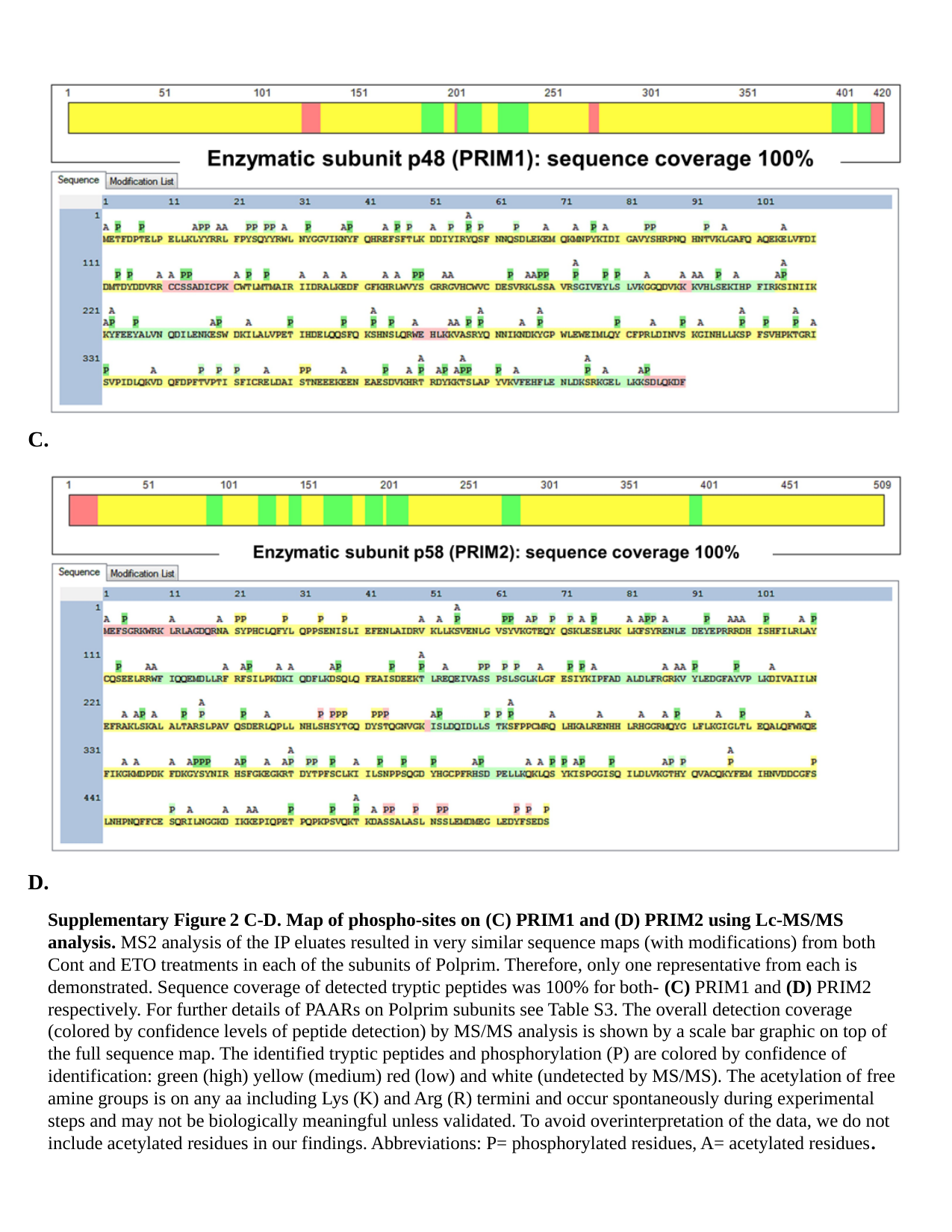

C.
D.
Supplementary Figure 2 C-D. Map of phospho-sites on (C) PRIM1 and (D) PRIM2 using Lc-MS/MS analysis. MS2 analysis of the IP eluates resulted in very similar sequence maps (with modifications) from both Cont and ETO treatments in each of the subunits of Polprim. Therefore, only one representative from each is demonstrated. Sequence coverage of detected tryptic peptides was 100% for both- (C) PRIM1 and (D) PRIM2 respectively. For further details of PAARs on Polprim subunits see Table S3. The overall detection coverage (colored by confidence levels of peptide detection) by MS/MS analysis is shown by a scale bar graphic on top of the full sequence map. The identified tryptic peptides and phosphorylation (P) are colored by confidence of identification: green (high) yellow (medium) red (low) and white (undetected by MS/MS). The acetylation of free amine groups is on any aa including Lys (K) and Arg (R) termini and occur spontaneously during experimental steps and may not be biologically meaningful unless validated. To avoid overinterpretation of the data, we do not include acetylated residues in our findings. Abbreviations: P= phosphorylated residues, A= acetylated residues.

### Slide 5
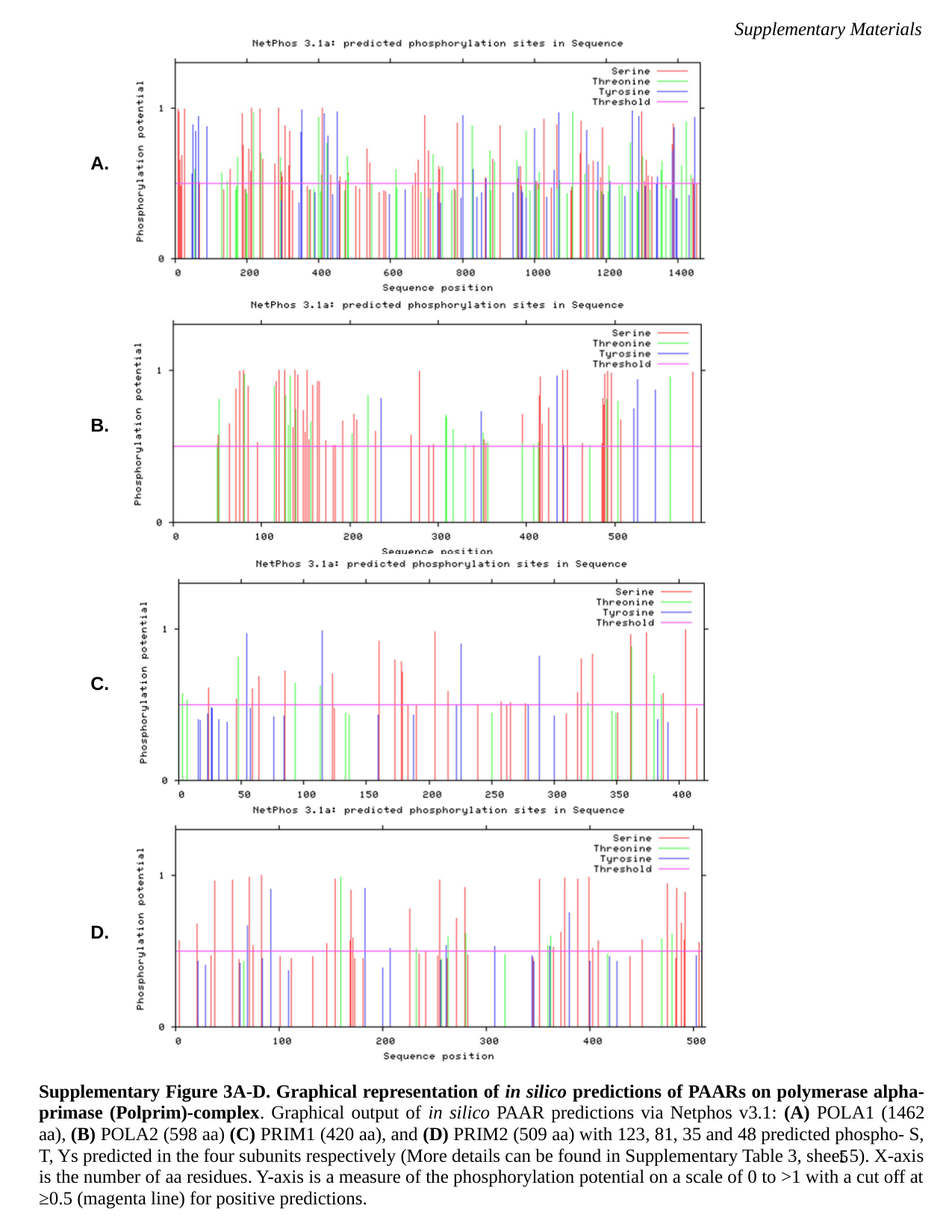

Supplementary Materials
A.
B.
C.
D.
Supplementary Figure 3A-D. Graphical representation of in silico predictions of PAARs on polymerase alpha-primase (Polprim)-complex. Graphical output of in silico PAAR predictions via Netphos v3.1: (A) POLA1 (1462 aa), (B) POLA2 (598 aa) (C) PRIM1 (420 aa), and (D) PRIM2 (509 aa) with 123, 81, 35 and 48 predicted phospho- S, T, Ys predicted in the four subunits respectively (More details can be found in Supplementary Table 3, sheet 5). X-axis is the number of aa residues. Y-axis is a measure of the phosphorylation potential on a scale of 0 to >1 with a cut off at ≥0.5 (magenta line) for positive predictions.
5

### Slide 6
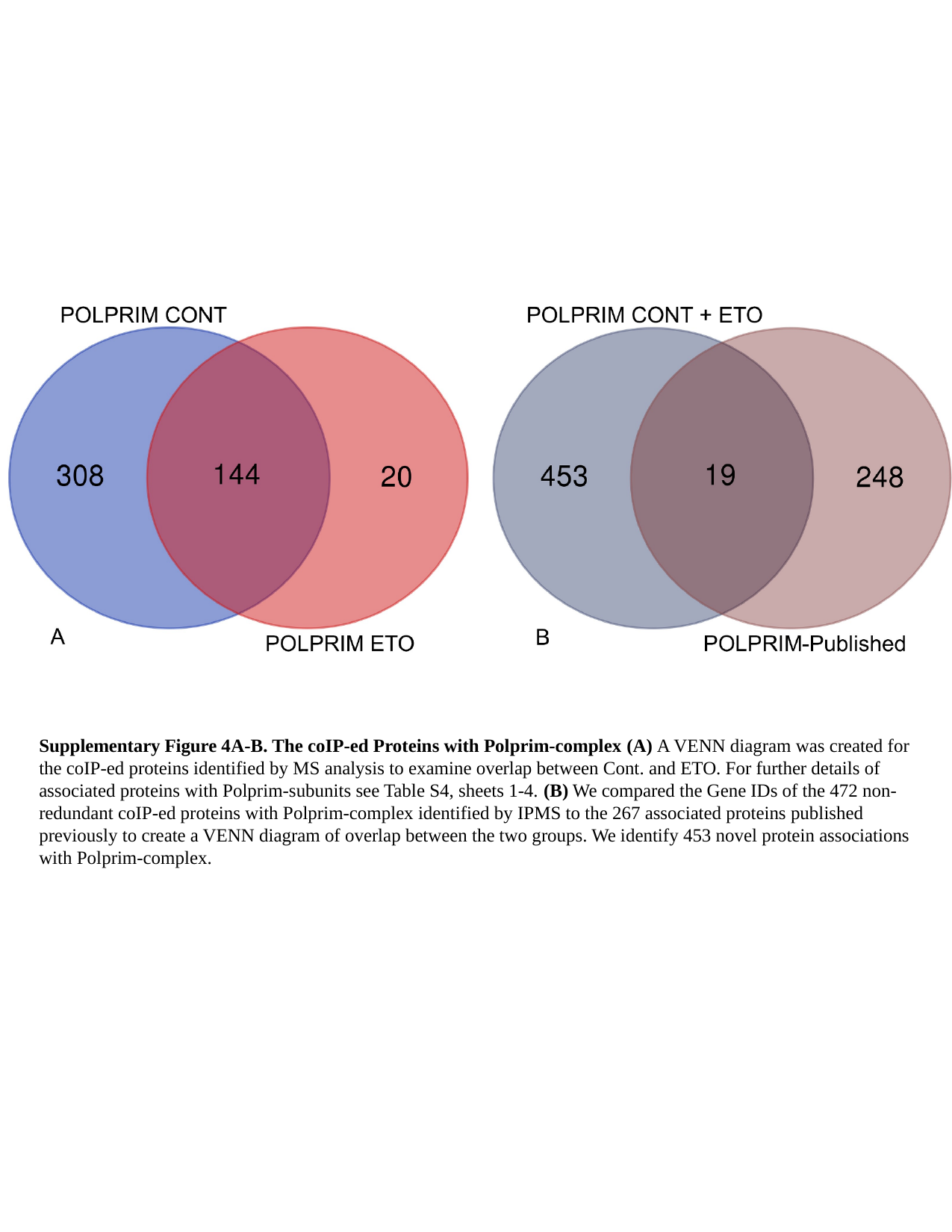

Supplementary Figure 4A-B. The coIP-ed Proteins with Polprim-complex (A) A VENN diagram was created for the coIP-ed proteins identified by MS analysis to examine overlap between Cont. and ETO. For further details of associated proteins with Polprim-subunits see Table S4, sheets 1-4. (B) We compared the Gene IDs of the 472 non-redundant coIP-ed proteins with Polprim-complex identified by IPMS to the 267 associated proteins published previously to create a VENN diagram of overlap between the two groups. We identify 453 novel protein associations with Polprim-complex.

### Slide 7
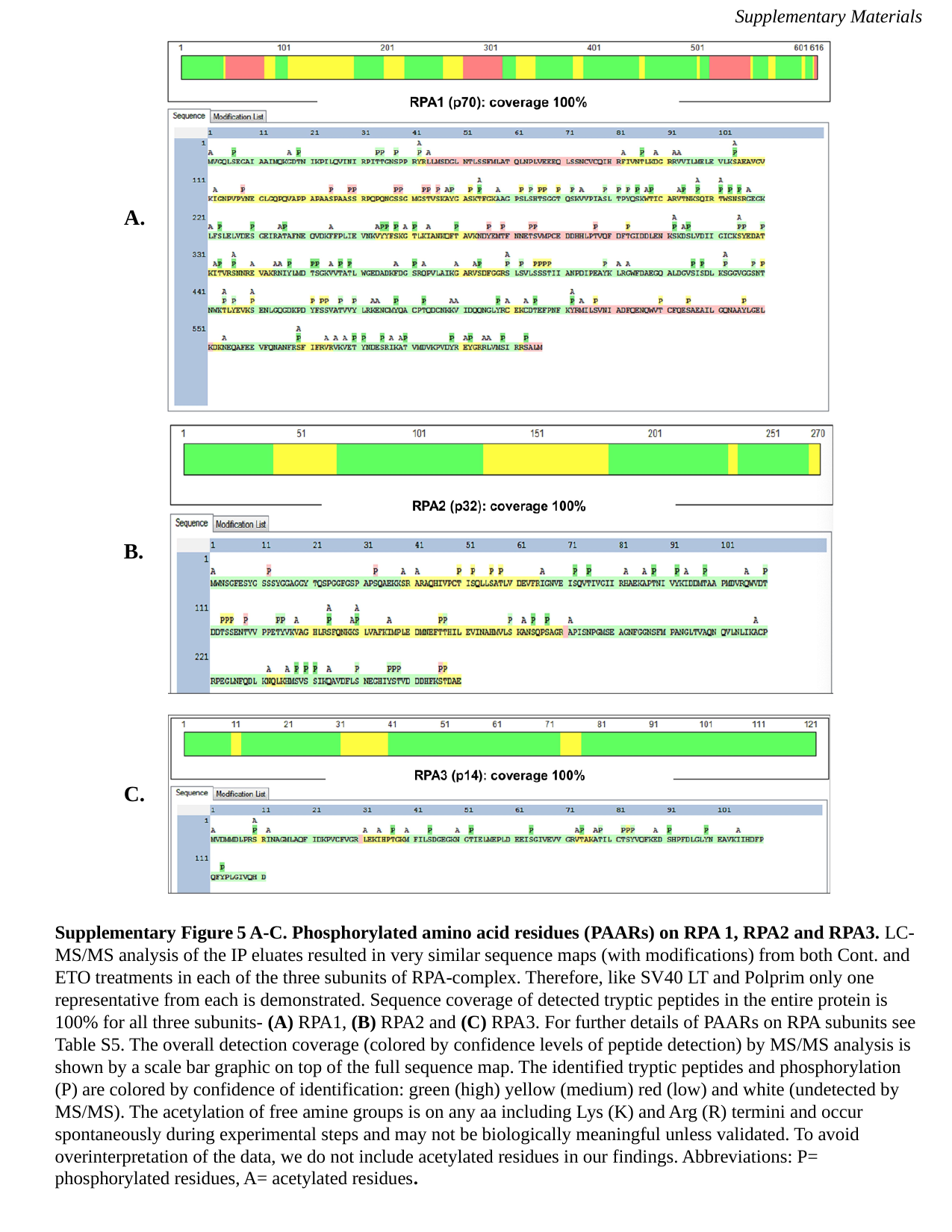

Supplementary Materials
A.
B.
C.
Supplementary Figure 5 A-C. Phosphorylated amino acid residues (PAARs) on RPA 1, RPA2 and RPA3. LC-MS/MS analysis of the IP eluates resulted in very similar sequence maps (with modifications) from both Cont. and ETO treatments in each of the three subunits of RPA-complex. Therefore, like SV40 LT and Polprim only one representative from each is demonstrated. Sequence coverage of detected tryptic peptides in the entire protein is 100% for all three subunits- (A) RPA1, (B) RPA2 and (C) RPA3. For further details of PAARs on RPA subunits see Table S5. The overall detection coverage (colored by confidence levels of peptide detection) by MS/MS analysis is shown by a scale bar graphic on top of the full sequence map. The identified tryptic peptides and phosphorylation (P) are colored by confidence of identification: green (high) yellow (medium) red (low) and white (undetected by MS/MS). The acetylation of free amine groups is on any aa including Lys (K) and Arg (R) termini and occur spontaneously during experimental steps and may not be biologically meaningful unless validated. To avoid overinterpretation of the data, we do not include acetylated residues in our findings. Abbreviations: P= phosphorylated residues, A= acetylated residues.

### Slide 8
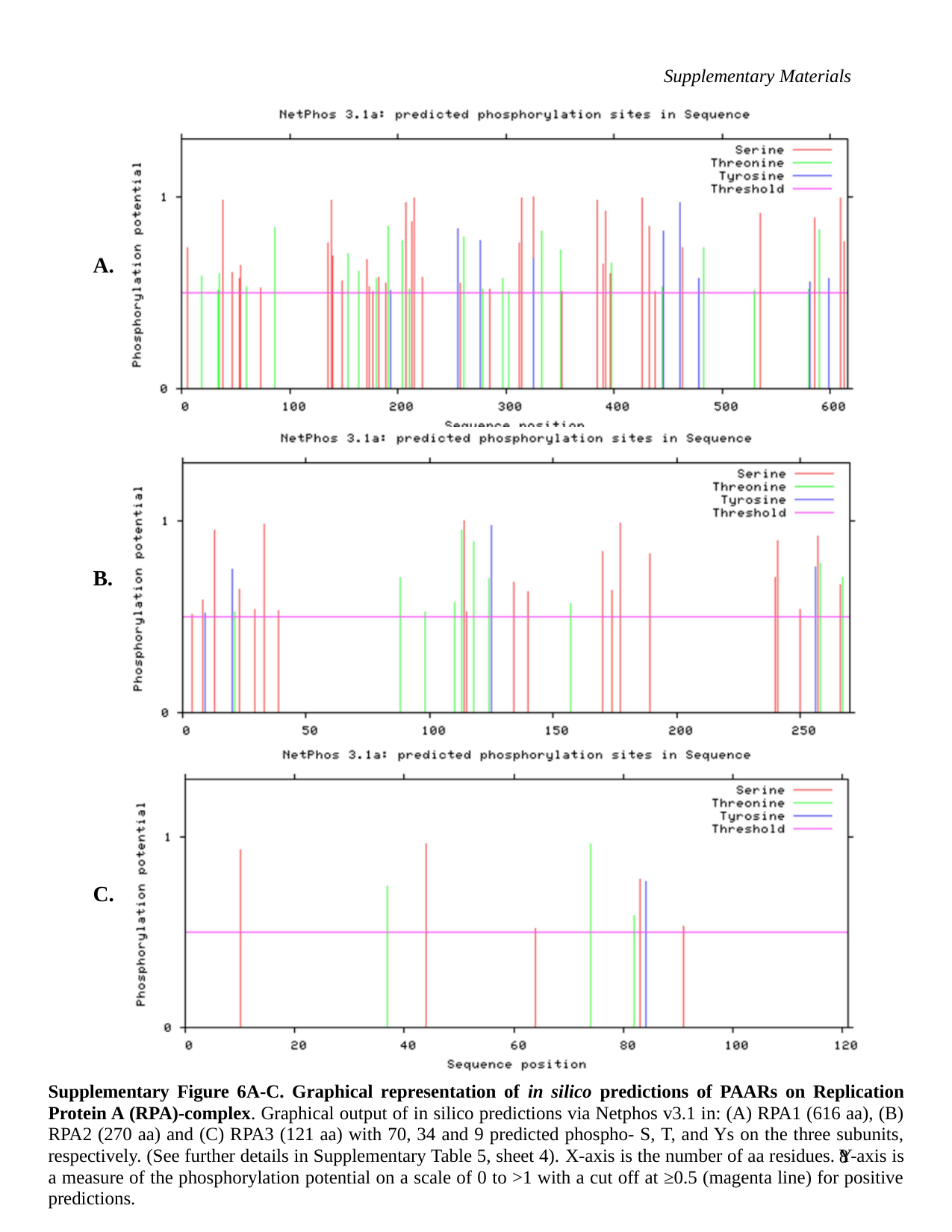

Supplementary Materials
A.
B.
C.
Supplementary Figure 6A-C. Graphical representation of in silico predictions of PAARs on Replication Protein A (RPA)-complex. Graphical output of in silico predictions via Netphos v3.1 in: (A) RPA1 (616 aa), (B) RPA2 (270 aa) and (C) RPA3 (121 aa) with 70, 34 and 9 predicted phospho- S, T, and Ys on the three subunits, respectively. (See further details in Supplementary Table 5, sheet 4). X-axis is the number of aa residues. Y-axis is a measure of the phosphorylation potential on a scale of 0 to >1 with a cut off at ≥0.5 (magenta line) for positive predictions.
8

### Slide 9
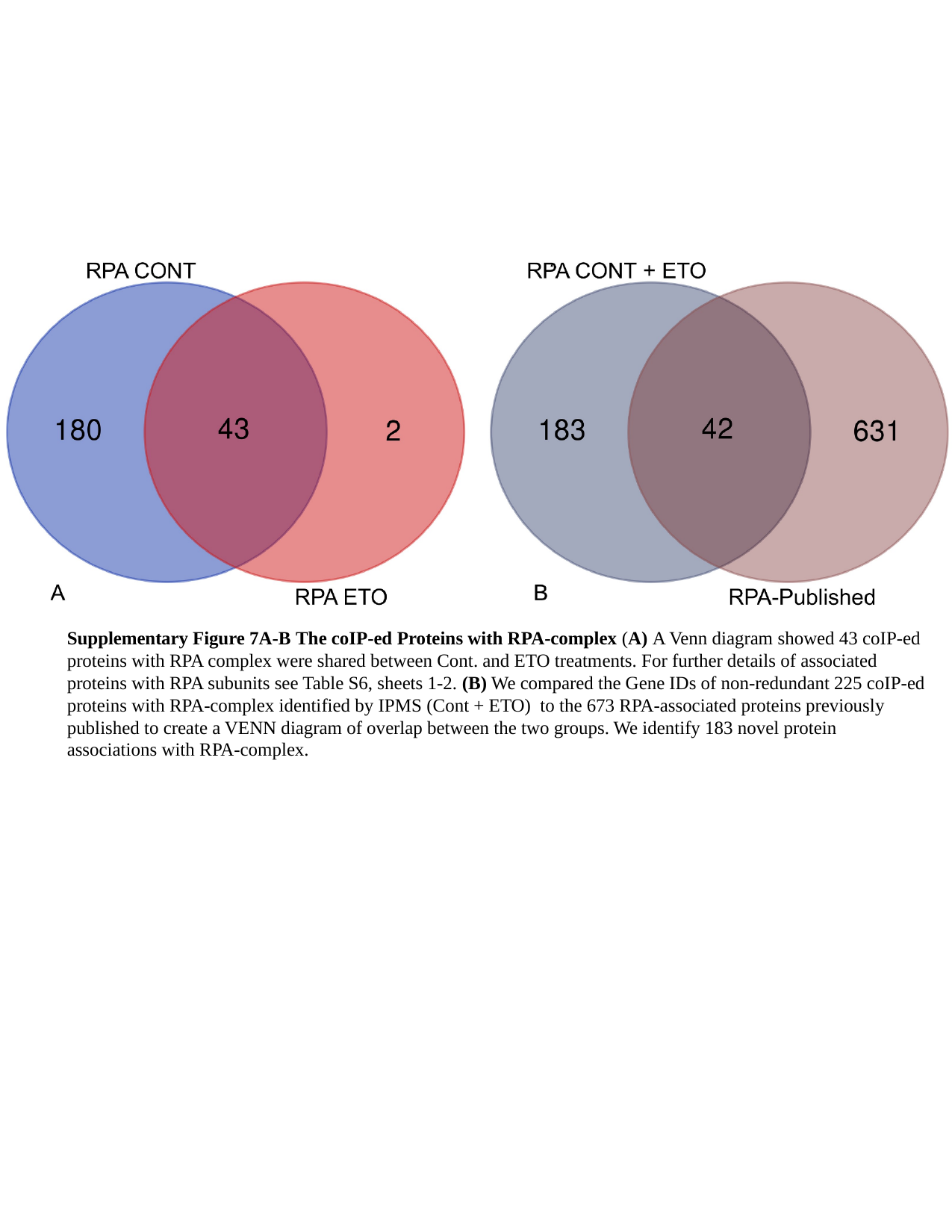

Supplementary Figure 7A-B The coIP-ed Proteins with RPA-complex (A) A Venn diagram showed 43 coIP-ed proteins with RPA complex were shared between Cont. and ETO treatments. For further details of associated proteins with RPA subunits see Table S6, sheets 1-2. (B) We compared the Gene IDs of non-redundant 225 coIP-ed proteins with RPA-complex identified by IPMS (Cont + ETO) to the 673 RPA-associated proteins previously published to create a VENN diagram of overlap between the two groups. We identify 183 novel protein associations with RPA-complex.

### Slide 10
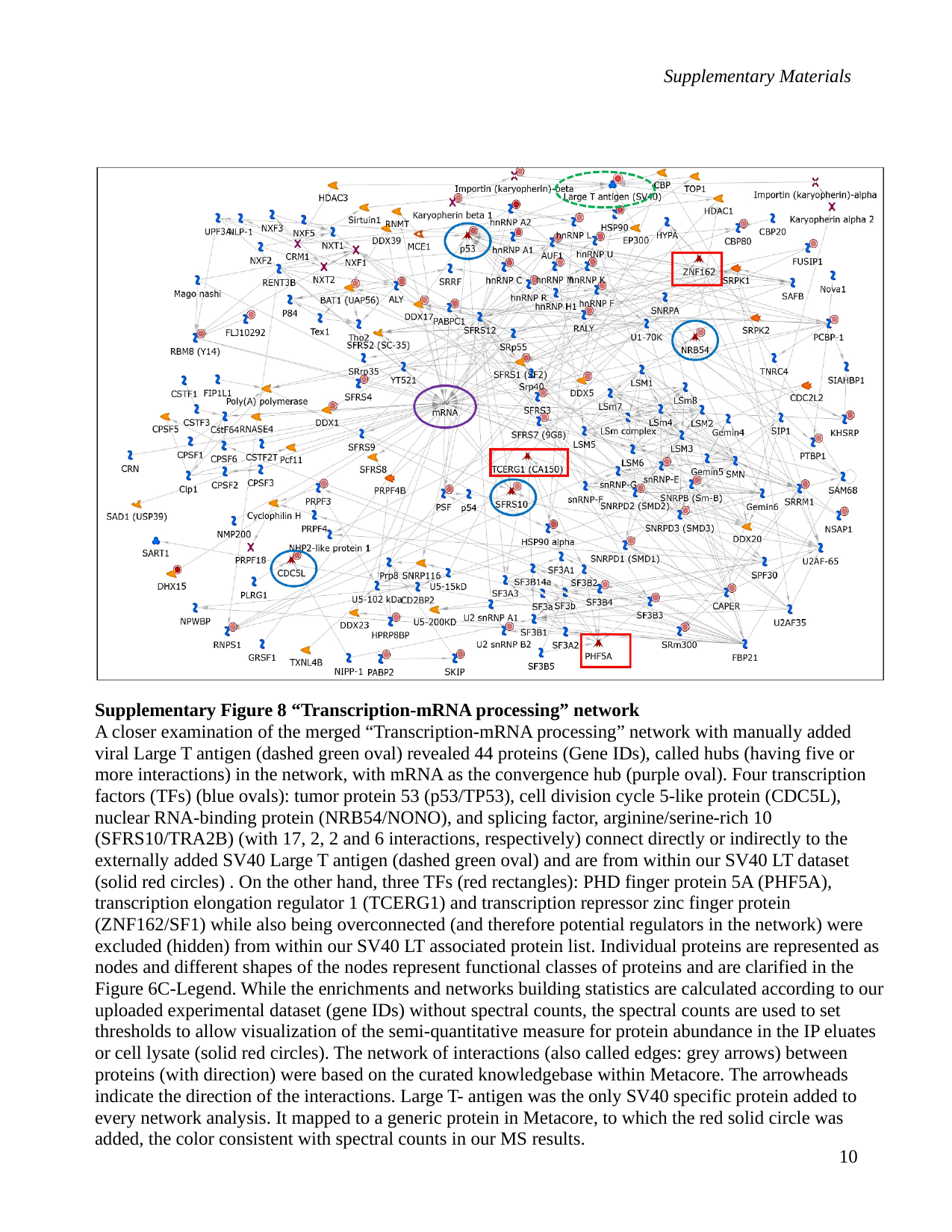

Supplementary Materials
Supplementary Figure 8 “Transcription-mRNA processing” network
A closer examination of the merged “Transcription-mRNA processing” network with manually added viral Large T antigen (dashed green oval) revealed 44 proteins (Gene IDs), called hubs (having five or more interactions) in the network, with mRNA as the convergence hub (purple oval). Four transcription factors (TFs) (blue ovals): tumor protein 53 (p53/TP53), cell division cycle 5-like protein (CDC5L), nuclear RNA-binding protein (NRB54/NONO), and splicing factor, arginine/serine-rich 10 (SFRS10/TRA2B) (with 17, 2, 2 and 6 interactions, respectively) connect directly or indirectly to the externally added SV40 Large T antigen (dashed green oval) and are from within our SV40 LT dataset (solid red circles) . On the other hand, three TFs (red rectangles): PHD finger protein 5A (PHF5A), transcription elongation regulator 1 (TCERG1) and transcription repressor zinc finger protein (ZNF162/SF1) while also being overconnected (and therefore potential regulators in the network) were excluded (hidden) from within our SV40 LT associated protein list. Individual proteins are represented as nodes and different shapes of the nodes represent functional classes of proteins and are clarified in the Figure 6C-Legend. While the enrichments and networks building statistics are calculated according to our uploaded experimental dataset (gene IDs) without spectral counts, the spectral counts are used to set thresholds to allow visualization of the semi-quantitative measure for protein abundance in the IP eluates or cell lysate (solid red circles). The network of interactions (also called edges: grey arrows) between proteins (with direction) were based on the curated knowledgebase within Metacore. The arrowheads indicate the direction of the interactions. Large T- antigen was the only SV40 specific protein added to every network analysis. It mapped to a generic protein in Metacore, to which the red solid circle was added, the color consistent with spectral counts in our MS results.
10

### Slide 11
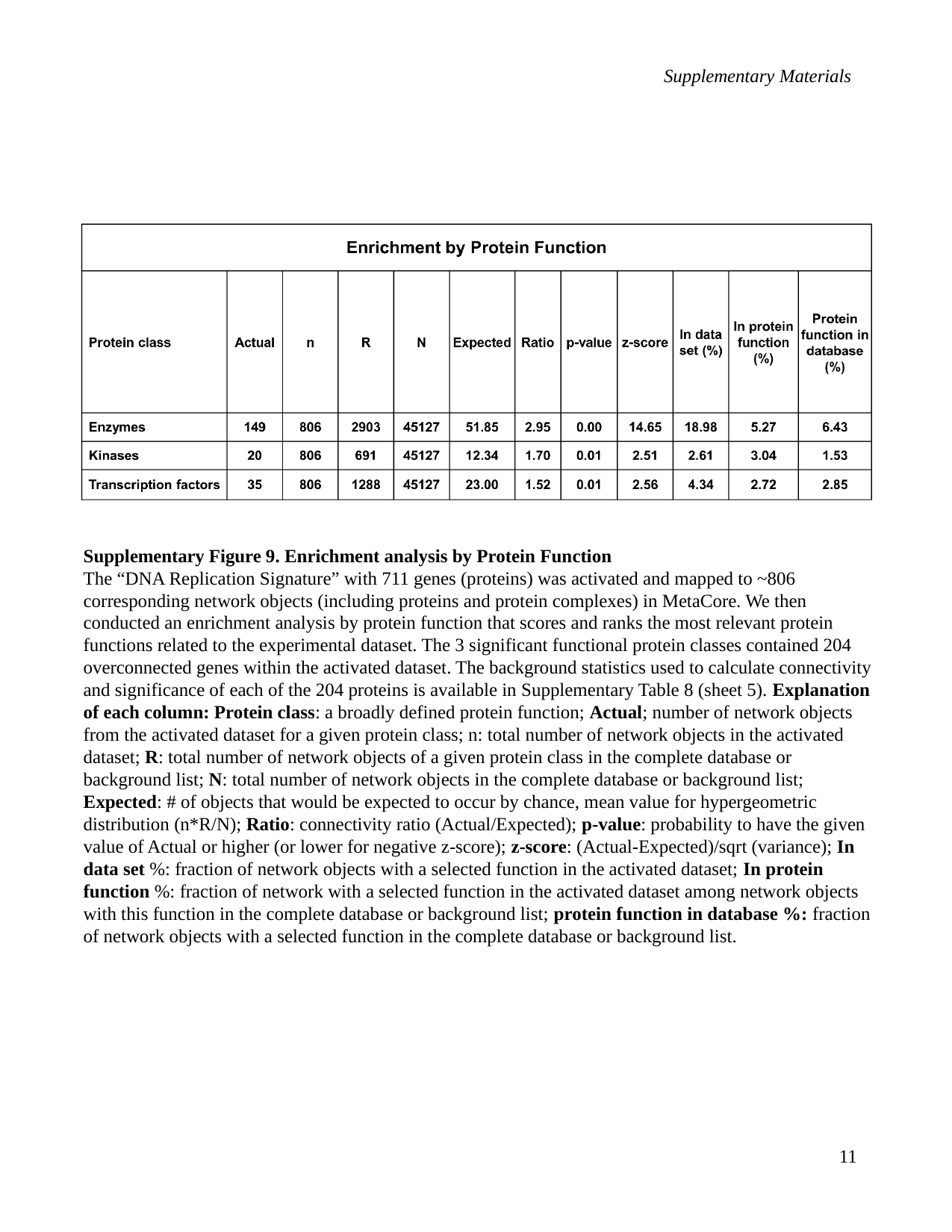

Supplementary Materials
Supplementary Figure 9. Enrichment analysis by Protein Function
The “DNA Replication Signature” with 711 genes (proteins) was activated and mapped to ~806 corresponding network objects (including proteins and protein complexes) in MetaCore. We then conducted an enrichment analysis by protein function that scores and ranks the most relevant protein functions related to the experimental dataset. The 3 significant functional protein classes contained 204 overconnected genes within the activated dataset. The background statistics used to calculate connectivity and significance of each of the 204 proteins is available in Supplementary Table 8 (sheet 5). Explanation of each column: Protein class: a broadly defined protein function; Actual; number of network objects from the activated dataset for a given protein class; n: total number of network objects in the activated dataset; R: total number of network objects of a given protein class in the complete database or background list; N: total number of network objects in the complete database or background list; Expected: # of objects that would be expected to occur by chance, mean value for hypergeometric distribution (n*R/N); Ratio: connectivity ratio (Actual/Expected); p-value: probability to have the given value of Actual or higher (or lower for negative z-score); z-score: (Actual-Expected)/sqrt (variance); In data set %: fraction of network objects with a selected function in the activated dataset; In protein function %: fraction of network with a selected function in the activated dataset among network objects with this function in the complete database or background list; protein function in database %: fraction of network objects with a selected function in the complete database or background list.
11

### Slide 12
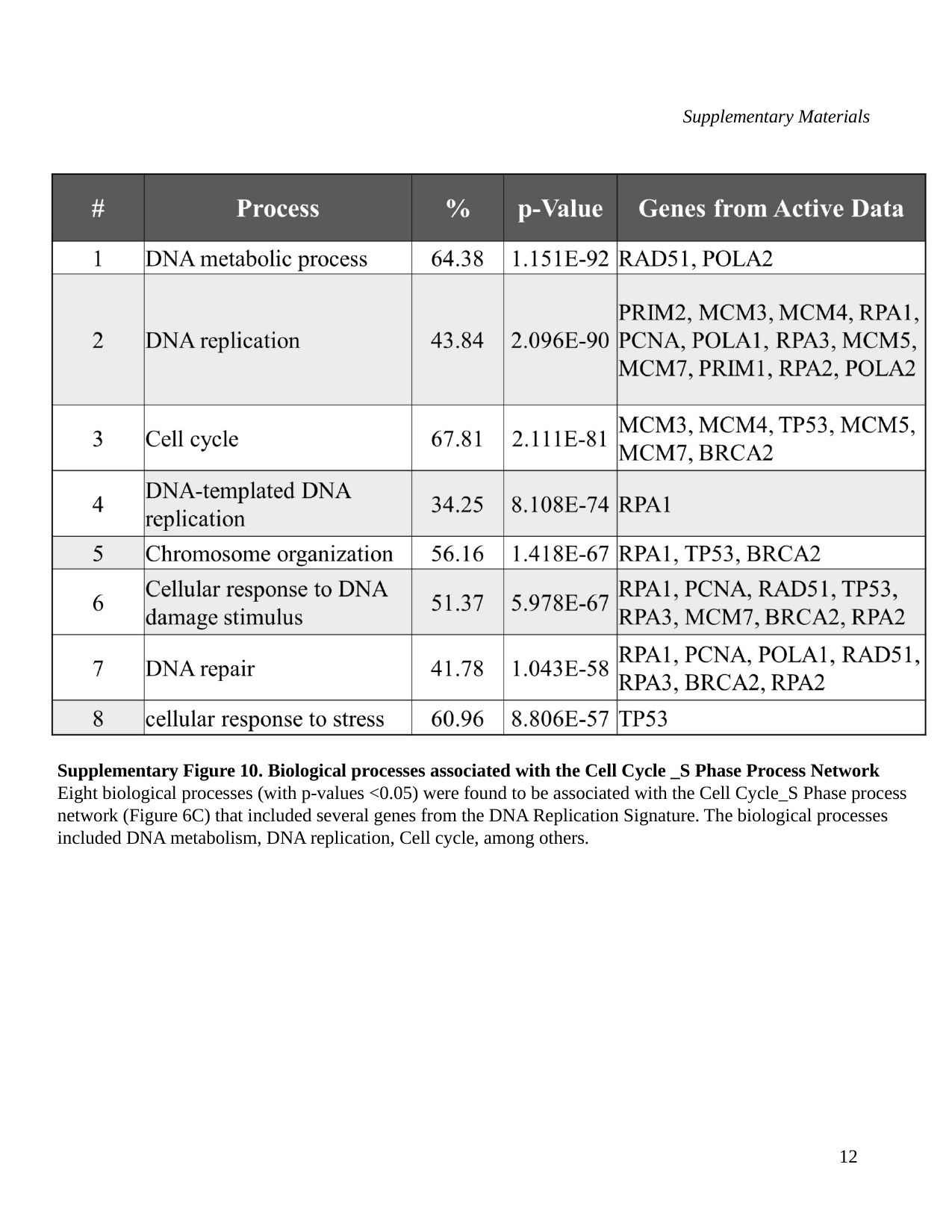

Supplementary Materials
Supplementary Figure 10. Biological processes associated with the Cell Cycle _S Phase Process Network Eight biological processes (with p-values <0.05) were found to be associated with the Cell Cycle_S Phase process network (Figure 6C) that included several genes from the DNA Replication Signature. The biological processes included DNA metabolism, DNA replication, Cell cycle, among others.
12
